# Genes and sites under adaptation at the phylogenetic scale also exhibit adaptation at the population-genetic scale

**DOI:** 10.1101/2022.09.23.509132

**Authors:** T. Latrille, N. Rodrigue, N. Lartillot

**Affiliations:** Université de Lyon, CNRS, LBBE UMR 5558, Villeurbanne, France; École Normale Supérieure de Lyon, Université de Lyon, Lyon, France; Department of Computational Biology, Université de Lausanne, Lausanne, Switzerland; Department of Biology, Institute of Biochemistry, and School of Mathematics and Statistics, Carleton University, Ottawa, Canada

**Keywords:** Adaptation, phylogenetic, population genetics, codon models

## Abstract

Adaptation in protein-coding sequences can be detected from multiple sequence alignments across species, or alternatively by leveraging polymorphism data inside a population. Across species, quantification of the adaptive rate relies on phylogenetic codon models, classically formulated in terms of the ratio of non-synonymous over synonymous substitution rates. Evidence of an accelerated non-synonymous substitution rate is considered a signature of pervasive adaptation. However, because of the background of purifying selection, these models are potentially limited in their sensitivity. Recent developments have led to more sophisticated mutation-selection codon models aimed at making a more detailed quantitative assessment of the interplay between mutation, purifying and positive selection. In this study, we conducted a large-scale exome-wide analysis of placental mammals with mutation-selection models, assessing their performance at detecting proteins and sites under adaptation. Importantly, mutation-selection codon models are based on a population-genetic formalism and thus are directly comparable to McDonald & Kreitman tests at the population level to quantify adaptation. Taking advantage of this relationship between phylogenetic and population genetics, we integrated divergence and polymorphism data across the entire exome for 29 populations across 7 genera, and showed that proteins and sites detected to be under adaptation at the phylogenetic scale are also under adaptation at the population-genetic scale. Altogether, our exome-wide analysis shows that phylogenetic mutation-selection codon models and population-genetic test of adaptation can be reconciled and are congruent, paving the way for integrative models and analyses across individuals and populations.

**Significance Statement:** Detecting genes under adaptation represents a key step in the decoding of genomes. Several methods have been proposed, focussing either on the short time scale (population genetics, e.g. human populations), or on the long time scale (phylogenetics, e.g. across mammals). However, the accuracy of these methods is still under debate, and it is still unclear whether the signatures of adaptation are congruent across evolutionary scales. In this study, using novel phylogenetic methods and gathering genome data across and within species, we show that the signatures of adaptation at the phylogenetic and population-genetic scales can be reconciled. While providing a mutual confirmation of the two approaches, our work paves the way for further methodological integration between micro- and macro-evolutionary genomics.

## Introduction

Present-day genetic sequences are informative of populations’ past evolutionary history and can carry signatures of selection at different scales. One main goal of molecular evolution is to disentangle and quantify the intensity of neutral, adaptive and purifying evolution acting on sequences, leveraging variations in sequences between and within species. Theoretically, in order to detect adaptive evolution, one must have data where part of the sequence is known to be under a neutral regime, which can be used as a null model. In the case of protein-coding DNA sequences, synonymous sites are usually taken as proxies for neutral sites, although there are instance where they are indeed under selection[1–3]. Non-synonymous mutations, on the other hand, might be under a mixture of varying degrees of adaptive and purifying selection. Contrasting synonymous and non-synonymous changes, two different types of methods have emerged to quantify both positive and purifying selection acting on protein-coding sequences. One method, stemming from phylogeny, uses a multiple sequence alignment comprised of genes from different species and relies on codon models to deduce the selective regime from the patterns of non-synonymous versus synonymous substitutions[4, 5]. Starting with the work of McDonald & Kreitman[6], another method, stemming from population genetics, contrasts polymorphism within a population and divergence to a closely related species.

At the population-genetic scale, one of the most widely used tests for adaptation relies on the substitutions between two closely related species and polymorphism within one population[6]. Under a strict neutral model (i.e. assuming non-synonymous mutations are either neutral or strongly selected), the ratios of non-synonymous polymorphisms over synonymous polymorphisms (*π*_*N*_ */π*_*S*_) and of non-synonymous substitutions over synonymous substitutions (*d*_*N*_ */d*_*S*_) are expected to be equal. If, in addition, strongly advantageous mutations occur, they are fixed rapidly in the population, thus contributing solely to divergence but not to polymorphism. As a result, the positive difference between *d*_*N*_ */d*_*S*_ and *π*_*N*_ */π*_*S*_ gives an estimate of the adaptive rate *ω*_A_ = *d*_*N*_ */d*_*S*_ *− π*_*N*_ */π*_*S*_[7]. This simple argument is not strictly valid in the presence of moderately deleterious non-synonymous mutations, which can segregate at substantial frequency in the population without reaching fixation, thus contributing solely to polymorphism, and not to divergence, potentially resulting on an underestimation of the rate of adaptive evolution[8]. Subsequent developments have tried to correct for this effect by relying on an explicit nearly-neutral model[9, 10], so as to estimate the rate of evolution expected in the absence of adaptation (called *ω*_0_) based on polymorphism, and then to compare it with the rate of evolution, *ω* = *d*_*N*_ */d*_*S*_, to get an estimate of the rate of adaptation as *ω*_A_ = *ω − ω*_0_.

In their current formulation, phylogeny-based methods rely on the ratio of non-synonymous substitutions over synonymous substitutions, called *ω*[4, 5]. Assuming synonymous mutations are neutral, *ω >* 1 signals an excess in the rate of non-synonymous substitutions compared to the neutral expectation, indicating that the protein is under adaptive evolution. Conversely, a deficit in non-synonymous substitutions, leading to *ω <* 1, means the protein is under purifying selection. In practice, proteins are typically under a mix of adaptive and purifying selection dominated by the latter, thus typically leading to an *ω <* 1 even in the presence of positive selection. At a finer scale, site models can detect a specific site (*i*) of the sequence with a *ω*^(*i*)^ *>* 1[11, 12]. Site models have the advantage of greater sensitivity and the ability to pinpoint where positive selection acts on the protein. However, even at the level of a single site under recurrent adaptation, not all amino-acids are expected to be adaptive, leading to *ω*^(*i*)^ capturing a mix of adaptive and purifying selection, reducing the sensitivity of test. An alternative approach to detect adaptation would be to rely on an explicit nearly-neutral model as the null against which to detect deviations, similarly to the McDonald & Kreitman test. Recent development in this direction, the so-called phylogenetic mutation-selection models provide a null model by estimating the fitness landscape over amino acid sequences, for each site of the sequence[13–15]. At the mutation-selection balance, the probability for a specific codon to be fixed in the population is proportional to its fitness, and a mutation from a high fitness amino acid towards a low fitness amino acid will have a small probability of fixation, genuinely accounting for purifying selection. Conversely, only nearly-neutral mutations between high fitness amino acids will tend to be permitted by the model, allowing for the explicit calculation of the nearly-neutral rate of non-synonymous substitutions at mutation-selection balance, called *ω*_0_[16, 17]. By contrasting *ω* estimated by *ω*-based codon models and *ω*_0_ calculated from mutation-selection models, one can hope to extract the rate of adaptation 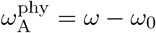.

Interestingly, the rate of adaptation is directly comparable between phylogenetic and population-genetic methods since both seek a deviation of *ω* from a nearly-neutral null model, estimated with mutation-selection models in phylogenetic context (*ω*_0_) or from standing polymorphism in a population-genetic context (*π*_*N*_ */π*_*S*_). This raises the question whether the two signals of adaptation are correlated, thus representing a unique opportunity to confront phylogeny-based and population-based methods. These two methods work over very different time scales, for that reason, they might be capturing different signals: long-term evolutionary Red-Queen for phylogeny-based methods versus events of adaptation in specific lineages for population-based methods. Nonetheless, we expect sites and proteins under long-term evolutionary Red-Queen regimes to maintain their signal of adaptation in several independent lineages for which the McDonald & Kreitman test is applied.

Accordingly, in this study, we first applied *ω*-based and mutation-selection codon models to whole exome data from placental mammals, so as to quantify the rate 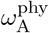 for each site and protein and detect signatures of adaptive evolution at the phylogenetic scale. Then, we developed a pipeline integrating (and aligning) divergence and polymorphism data across the entire exome for 29 populations across 7 genera, namely *Equus, Canis, Bos, Capra, Ovis, Chlorocebus* and *Homo*. Finally, using this pipeline, we assessed the congruence between the phylogeny-based and population-based approaches, by testing if the group of sequences detected with a high rate of adaptation in the phylogeny-based method also displays a high rate of adaptation according to the population-based method.

## Results

### Detecting genes and sites under adaptation

We derived a two-step approach (see methods), which we applied to a set of alignments of orthologous genes at the scale of placental mammals. The *d*_*N*_ */d*_*S*_ estimated by the site model (*ω*) is plotted against the *d*_*N*_ */d*_*S*_ predicted by the nearly-neutral mutation-selection model (*ω*_0_) for genes (scatter plot in fig. 1A) and sites (density plot in fig. 1B). An excess of *ω* relative to *ω*_0_ is a typical signature of ongoing positive selection[17, 18]. Accordingly, genes, or sites, were considered to be under an adaptive regime (in red) if the value of their *ω* is higher than that of their *ω*_0_, with non-overlapping 95% posterior credibility intervals. This procedure retrieved 822 out 14,509 genes, which are putatively under a long-term evolutionary Red-Queen regime. At the site level, the nearly-neutral assumption appears to be rejected for 104,129 out of 8,895,374 sites. Of note, this selection procedure is not meant as a routine statistical test, but only as an enrichment procedure, for the needs of the subsequent analysis shown below. In practice, this selection is likely to be conservative and to have a rate of false discovery of the order of 1% at the gene-level, and 5% at the site-level (see methods).

**Figure 1:**
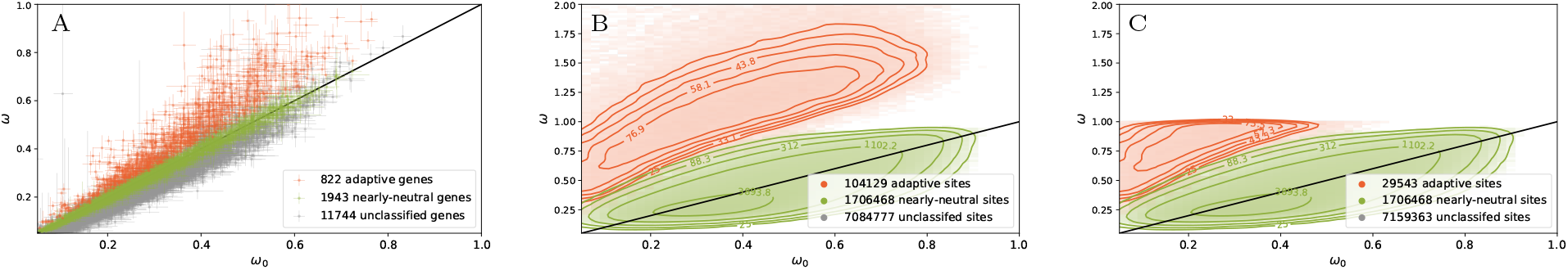
Detection of protein-coding sequences ongoing adaptation at the phylogenetic scale. *ω* estimated by the site model against *ω*_0_ calculated by the mutation-selection model. Scatter plot of 14,509 genes in panel A, with 95% bayesian credible interval (*α* = 0.05). Density plot of sites in panel B and C. Genes or sites are then classified whether they detected as adaptive (*ω > ω*_0_ in red) or nearly-neutral (*ω ≃ ω*_0_ in green). In panel C, the set of sites detected exclusively by mutation-selection codon models have a mean *ω <* 1.

Of note, selection based on *ω > ω*_0_ is more sensitive than based on the commonly used criterion of *ω >* 1, since *ω*_0_ is always lower than 1 by definition[16]. Thus, we can uncover sites under adaptation (*ω > ω*_0_) with a mean *ω* lower than 1 (29,543 sites in fig. 1C). These sites could not have been detected by *ω*-based codon models relying on the criterion that *ω >* 1. At the gene level, only two genes have an estimated *ω >* 1, such that this distinction is not relevant.

### Ontology enrichment tests

Next, we investigated whether the genes classified as adaptive (*ω > ω*_0_) showed enrichment in specific ontology terms. Thus, we performed 775 instances of Fisher’s exact test to estimate ontology enrichment by contrasting with genes in the control group, not classified as adaptive. 42 ontologies are observed with a p-value (*p*_v_) corrected for multiple comparison (Holm–Bonferroni correction, 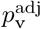) lower than the risk *α* = 0.05 (see table S1). At a finer scale, we weighted genes by their proportion of sites considered under adaptation with a *ω*-based site model (*ω >* 1, see table S2) or with a mutation-selection model (*ω > ω*_0_, see table S3). For each ontology, the proportion of sites under adaptation is compared between the set of genes sharing this given ontology and the rest of the genes (Mann-Whitney U test). The statistical test based on the the first criterion (*ω >* 1) is correlated with ontologies related to immune processes, while the statistical test based on the the second criterion (*ω > ω*_0_) is also correlated with ontologies related to the external membrane and cellular adhesion.

### Congruence between phylogeny- and population-based methods

Finally, we investigated whether the phylogeny-based and the population-based methods give congruent results in terms of detection of adaptive evolution (fig. 2). To do so, population genomic data were collected for 29 populations across 7 genera. For each population, *ω*_A_ as proposed by McDonald & Kreitman (MK)[6] was computed on the concatenate of the 822 genes classified as adaptive by the phylogeny-based method (red dots in fig. 2 and 3). This result was compared to a null distribution obtained by computing *ω*_A_ over sets of 822 genes that were randomly sampled (1,000 replicates) among the genes classifed as nearly-neutral according to the mutation-selection model (green violins in fig. 2 and 3). Importantly, the terminal lineages over which the population-genetic method was applied were not included in the phylogenetic analysis. As a result, the two methods are working on entirely non-overlapping compartments of the evolutionary history across mammals. For all 29 populations, the *ω*_A_ estimated by the population-genetic method was significantly higher for the putatively adaptive gene-set than for the putatively nearly-neutral gene sets of the same size (at a risk *α* = 0.05 corrected for multiple testing, Holm-Bonferroni correction). There is thus a good qualitative agreement between the two methods as to what they capture and interpret as positive selection at the gene level.

**Figure 2:**
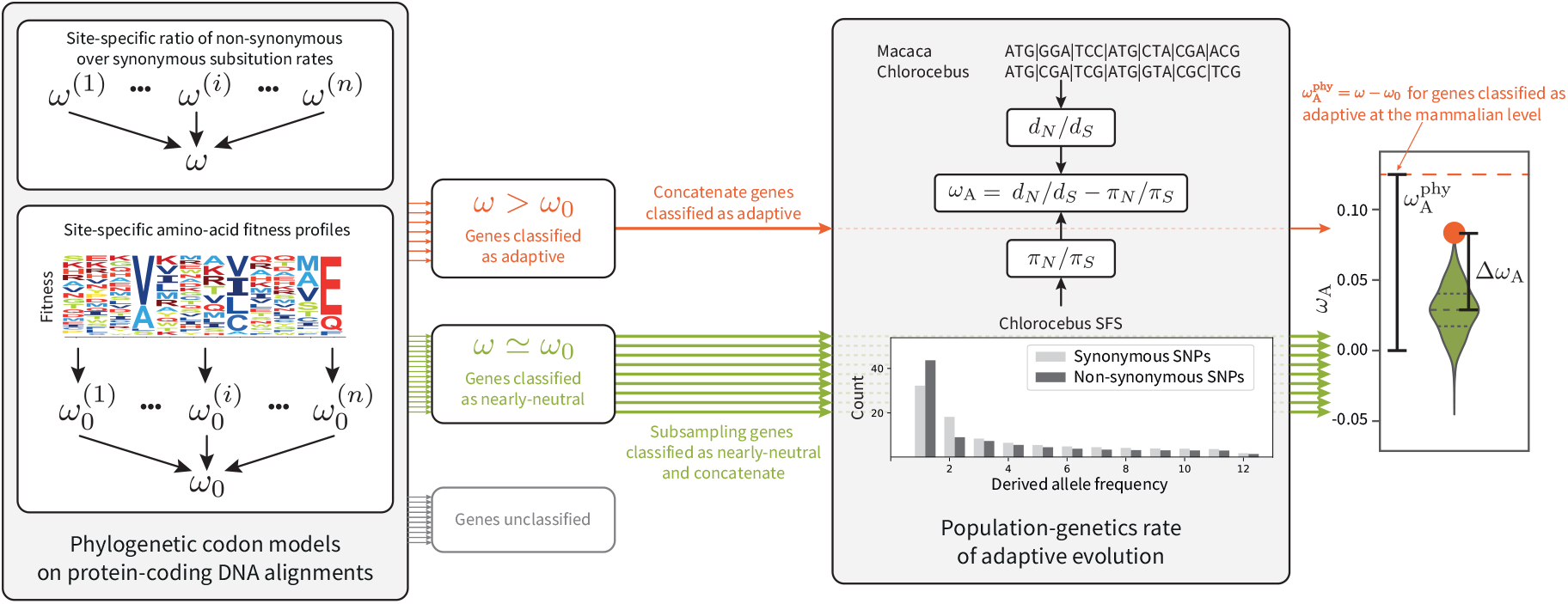
Integrating divergence and polymorphism for the detection of adaptation. At the phylogenetic level, *ω* (classical codon models) and *ω*_0_ (mutation-selection codon models) are computed from protein-coding DNA alignments, allowing to classify genes into adaptive (in red) and nearly-neutral (in green) regime. At the population-genetic level, for each population, *ω*_A_ is computed on the concatenate of genes classified as under adaptation. The result is compared to the empirical null distribution of *ω*_A_ in each population, obtained by randomly sampling (1,000 replicates) a subset under a nearly-neutral regime.

**Figure 3:**
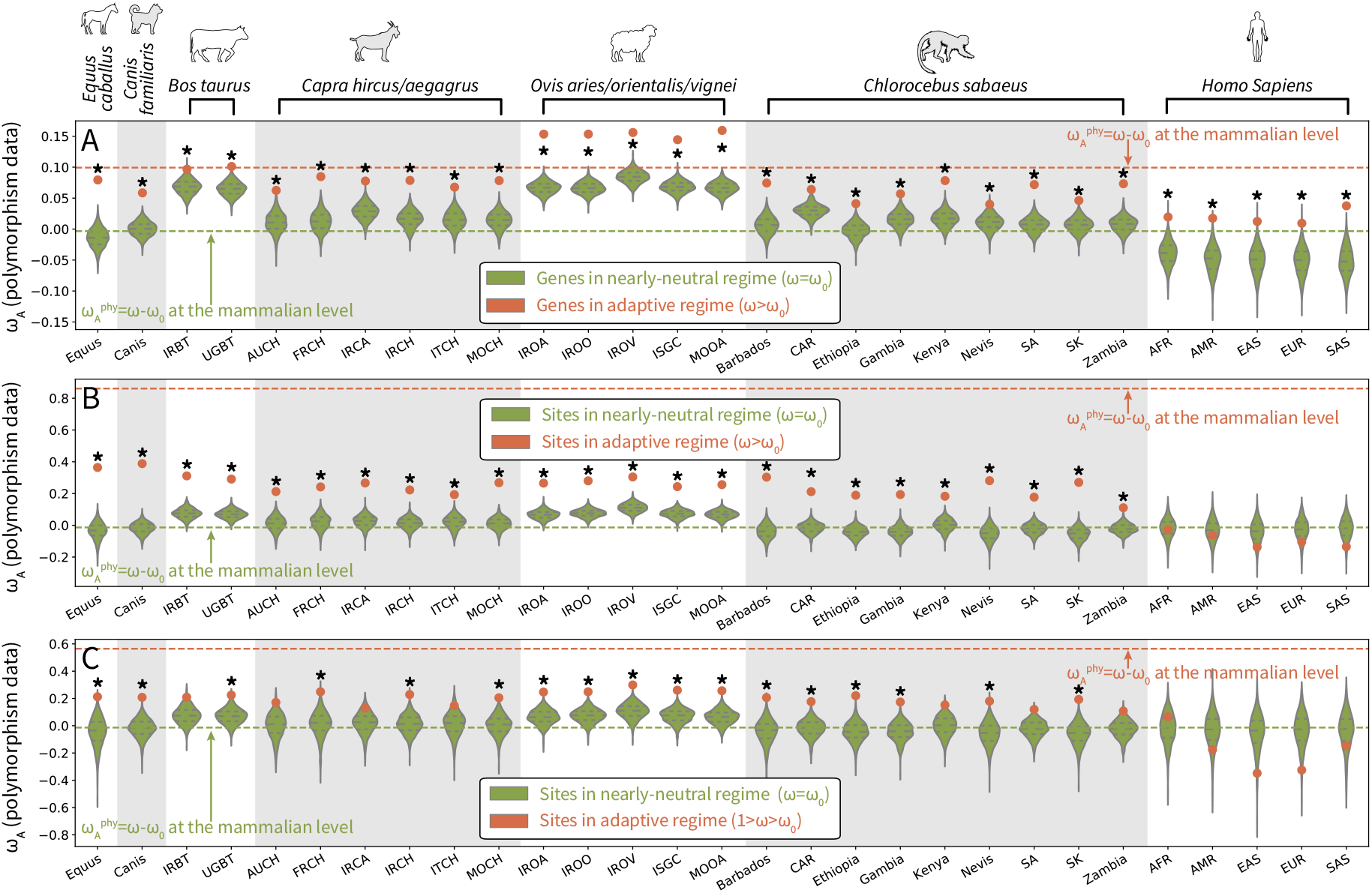
Enrichment of adaptation at the population-genetic scale for 29 populations across 7 genera at the gene (panel A) and site (panels B and C) level. For each population, *ω*_A_ is computed on 822 genes (A) and 104,129 sites (B) having a high rate of adaptation at the phylogenetic scale (*ω > ω*_0_ in red). In panel C, the set of 29,543 sites are detected exclusively by mutation-selection codon models with a mean *ω <* 1. The result is compared to the empirical null distribution of *ω*_A_, obtained by randomly sampling (1,000 replicates) a subset of genes and sites under a nearly-neutral regime (violin plot in green). ^*∗*^ signify that the *p*_v_ corrected for multiple comparison (Holm–Bonferroni correction) is lower than the risk *α* = 0.05. The acronym of populations, and the quantitative value of *ω*_A_ and *p*_v_ are shown in table 1

The same procedure was applied at a finer scale with sites instead of genes. For each population, *ω*_A_ was computed on the concatenate of the 104,129 sites classified as adaptive by the phylogeny-based method, and compared to the empirical null distribution (fig. 3B) and table 1. Out of 29 populations, 24 have an *ω*_A_ estimated by the population-genetic method significantly higher for the putatively adaptive site-set than for the putatively nearly-neutral site-sets of the same size taken at random (at a risk *α* = 0.05 corrected for multiple testing, Holm-Bonferroni correction). Of note, the 5 populations for which the test is not significant are the human populations.

**Table 1:**
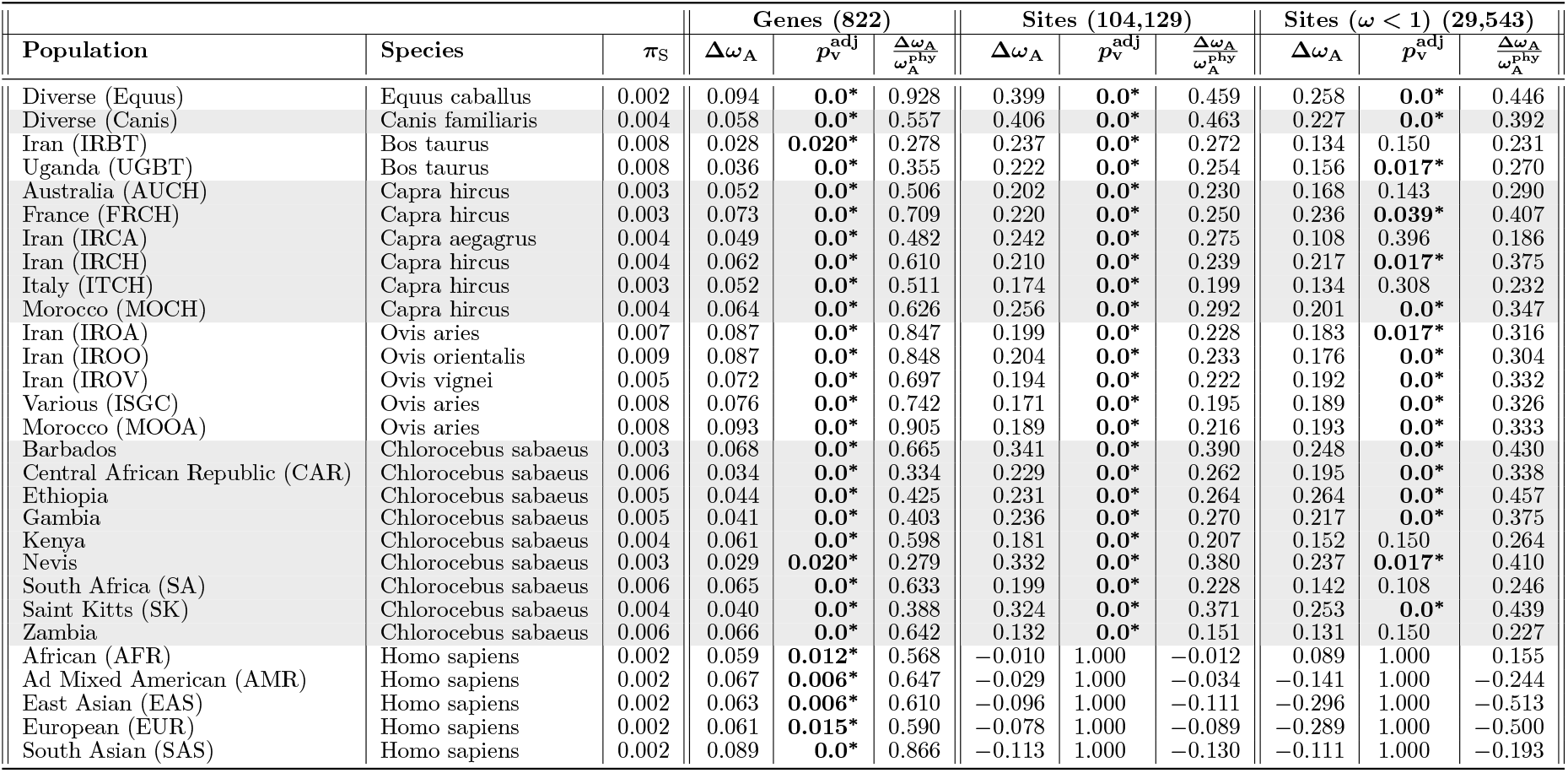
Across 29 populations (rows), table of quantitative value of Δ*ω*_A_ between the set classified as adaptive and nearly-neutral shown in fig. 3. 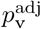 associated to the test are corrected for multiple comparison (Holm–Bonferroni correction, ^*∗*^ for 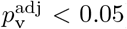). 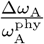 is the ratio of Δ*ω*_A_ at the population-genetic level and the phylogenetic level. *π*_S_ is the observed genetic diversity (number of SNPs per site) counted over synonymous sites.

Except for *Equus* and *Humans*, on average, the *ω*_A_ returned by MK is positive even for the putatively nearly-neutral replicates, and significantly so for *Bos* (*ω*_A_ in the range 0.65 *−* 0.68 for genes and site) and *Ovis* (*ω*_A_ in the range 0.66 *−* 0.84 for genes and sites). This suggests the presence of a background of positive selection captured by MK methods but not by phylogenetic methods. This background signal could correspond either to adaptation specifically present in the terminal lineages on which the MK method is applied and absent over the rest of the mammalian tree, or to low-intensity recurrent positive selection, present over the tree but nevertheless missed by phylogenetic methods, owing to a lack of sensitivity. Alternatively, part of it could be an artifact of MK methods, due for example to a recent demographic expansion (*Bos* and *Ovis* are the two among those analysed by the population-genetic approach showing the highest levels of synonymous diversity), or to a more general mismatch between short- and long-term effective population size (*N*_e_)[19].

Regardless of its exact cause, subtracting this background, so as to compare, not directly the *ω*_A_ of the population-genetic method, but the Δ*ω*_A_ between the putatively adaptive set and the control replicates, to the 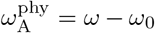 returned by the phylogenetic method, may give a more meaningful basis for a quantitative comparison between phylogenetic and population-genetic approaches (fig. 2). Of note, across all analyses shown in fig. 3A and B, this population-genetic Δ*ω*_A_ is always smaller than the phylogenetic 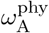. This asymmetry is expected, as a result of a selection bias: the genes of the test set were selected precisely for their high phylogenetic signal, while keeping a blind eye to their population-genetic signal. From this perspective, the ratio 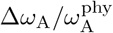can be interpreted as an estimate of the fraction of the total signal captured by the phylogenetic enrichment procedure that is confirmed by MK statistics. This ratio, hereafter called the confirmation rate, is indicated in table 1.

At the gene level, the confirmation rate is relatively high, ranging from 30% to up to 90%. At the site level, the confirmation rate is lower (30% on average), which could betray a higher rate of false discovery at the site level, or could be the result of subtle molecular evolutionary processes, such as intermittent adaptation (on some but not on all branches) or within-gene turnover (ongoing adaptation targeting different sites on different branches).

After discarding sites with a mean *ω >* 1, the remaining 29,543 sites classified as being under an adaptive regime have 1 *> ω > ω*_0_ and are specifically discovered by the mutation-selection approach. Since their *ω* is less than 1, they could not be detected by classical codon models. This raises the question of the empirical value of these findings. Indeed, while mutation-selection methods are more sophisticated and may therefore have a greater sensitivity, they may also be more prone to producing false positives. The phylogenetic/population-genetic confrontation developed here can be used to assess this important point. As shown in (fig. 3C) and table 1, out of 29 populations, for 17 out of the 29 populations that have been analysed, the confirmation rate is significantly positive (*α*=0.05, Holm–Bonferroni correction), and of the order of 10% on average. This importantly suggests the presence of a background of low-intensity positive selection, which is missed by classical codon models, but partially detected by mutation-selection models. In other words, the approach can detect a long-term evolutionary Red-Queen even for a site with *ω <* 1 that is still under adaptation at the population-genetic scale.

Because genes and sites classified as adaptive have a higher *ω* than genes/sites classified as nearly-neutral, *ω*_A_ could simply be higher for genes with higher *ω* due to this confounding factor. Thus we performed additional experiments where *ω* is controlled to be the same in the nearly-neutral replicate and the adaptive set of genes (fig. S2-6 and tables S5-9). Additionally, we performed the same experiments with a more stringent risk *α* = 0.005 (10 times greater) to classify genes and sites as adaptive (fig. S7-9 and tables S9-10). Our result are robust to both controlling for *ω* and with a different threshold to classify genes and sites as adaptive. Finally, we computed *ω*_A_ using the software polyDFE[20], which relies on the synonymous and non-synonymous unfolded site-frequency spectra (SFS) to estimate the distribution of fitness effects of mutations (DFE), and the rate of adaptation (fig. S10-17 and tables S11-18). Depending on the underlying assumptions for the shape of the DFE and the definition of *ω*_A_, we observed a wide range of *ω*_A_ both for the set of adaptive and nearly-neutral genes/sites. However, the statistical test for the enrichment of *ω*_A_ between the set of adaptive and nearly-neutral genes/sites gives results in the same direction whether computed by polyDFE or as McDonald & Kreitman [6] statistic, although the confirmation rate and the associated *p*_v_ are different.

## Discussion

Quantifying the rate of adaptation assumes that we can measure the rate of evolution and more importantly its deviation from a null model of evolution disallowing adaptation. For phylogenetic codon models, this null model of evolution is usually assumed to be neutral evolution and the rate of evolution computed as the ratio of non-synonymous over synonymous substitution rates (*ω*) is thus compared to 1. We first showed that, at the phylogenetic scale, *ω* can be compared to it’s expectation under the mutation-selection model (*ω*_0_), a nearly-neutral model instead of a neutral model of evolution, giving a quantitative estimate of the rate of adaptation as 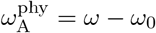. The application of this approach exome-wide across placental mammals suggests that 822 out of 14,509 proteins are under a long-term evolutionary Red-Queen, with ontology terms related to immune processes and the external membrane of cells. Enrichment of ontologies related to immune processes is expected, as found by many studies[12, 21, 22]. However, we also detect an enrichment with ontologies related to the external membrane and cell adhesion, which are the target of virus and parasites. Altogether, the mutation-selection method effectively detects adaptation regardless of the background of purifying selection, and returns reasonable candidates for adaptive evolution. Of note, in its current implementation, and unlike classical codon models[23, 24], the mutation-selection approach does not yet provide a proper and well-calibrated statistical test for calling genes or sites under adaptation with a well-controlled frequentist risk. This was not a problem in the enrichment analysis conducted in this article, which relies on downstream controls based on random permutations. Nevertheless, the encouraging results obtained here give a motivation for developing such a test, which should then have an increased power to detect adaptation, compared to classical codon models relying on the *ω >* 1 criterion.

At the population-genetic scale, the availability of approaches to detect adaptation[6, 25] raises the question whether the rate of adaptation calculated at the phylogenetic scale as 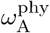 is congruent with the rate calculated at the population genetics scale by McDonald & Kreitman (MK)[6] as *ω*_A_ = *d*_*N*_ */d*_*S*_ *− π*_*N*_ */π*_*S*_. In this light, the set of genes and sites detected to be under adaptation at the phylogenetic scale showed a significant increase in *ω*_A_ such as inferred by population-based method (29 populations across 7 genera). Quantitatively, about 30% to 90% of the signal detected by the phylogeny-based approach is confirmed by the population-based approach. This result is in stark contrast with studies comparing *ω*-based codon models at the gene level with MK methods, which found that the set of genes detected at different scales does not seem to overlap beyond random expectations[26]. The reasons for this discrepancy are not totally clear. The use of different codon modeling strategies could play a role here. More fundamentally however, our study relies on a large and densely sampled phylogeny with ≃ 100 taxa across placental mammals, versus 5 *Drosophila* and 5 *Brassicaceae* in Chen *et al*. [26]. As a result, the phylogenetic aspect of our analysis benefits from an increased power, while being also inherently more focussed on genes characterized by recurrent adaptation over a very large evolutionary scale (i.e. long-term evolutionary Red Queens), for which population-genetic signals of adaptation may be more easily recovered. We thus showed empirically that the mutation-selection codon model provides a null (nearly-neutral) model from which we can disentangle purifying and adaptive evolution. However, our procedure still has some limitations.

Mutation-selection codon models assume a constant effective population size while it has been established that its fluctuations has a major effect on selection dynamics[27, 28]. Estimating changes in effective population size in a mutation-selection framework is possible[29], although too computational intensive in its current implementation to be performed genome-wide. Second, epistasis is not modeled while it can have a large effect on the response of the rate of evolution with change in population size[30]. More generally, pervasive epistasis generates an entrenchment of the amino acids[31–33], resulting in a slowing down of the rate evolution[17, 34] or a standstill[35]. Consequently, our estimation of the predicted rate of evolution computed at mutation-selection balance (*ω*_0_) is over-estimated given that epistasis is not taken into account, such that 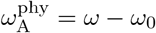is thus under-estimated. Altogether, we argue that our estimate of 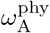 is conservative and could be increased by modeling epistasis (altough indirectly) within the mutation-selection framework[33].

On the other hand, at the population-genetic scale, the greatest limitation to detecting adaptation is the lack of power determined by the genetic diversity since polymorphisms are rare and estimation of *π*_*N*_ */π*_*S*_ requires to pool many sites for which SNPs are available. Since the effects of mild purifying selection are more pronounced on longer time scales (i.e. mildly deleterious mutations contribute disproportionately to polymorphism, compared to divergence), *ω*_A_ as computed by MK can be biased by moderately deleterious mutations[8, 36] and by the change in population size through time[37]. To overcome this bias, model-based approaches relying on the synonymous and non-synonymous site-frequency spectra (SFS) to estimate the distribution of fitness effects of mutations (DFE), so as to account for the contribution of mild selective effects to standing polymorphism, have been developed[9, 20] and are often used[10, 38]. However, the broad range of *ω*_A_ estimated on sets of genes/sites classified as nearly-neutral suggests that these models are lacking power, even more than the MK statistic, because of the sparsity of the SFS. Beside changes in population size biasing the estimation[19], we argue that inferring *ω*_A_ using an underlying DFE model is also highly sensitive to assumptions for the shape of the DFE and the definition of *ω*_A_. For example, the value of *ω*_A_ is computed as an integral, where the bounds of this integral is debated by different authors[10, 39]. It is thus relatively easy to change the definition of *ω*_A_ (fig. S12-15 and tables S13-16) or to constrain the underlying DFE (fig. S12-17 and tables S13-18) to obtain a wide range of *ω*_A_ on the same dataset. Taken together, we argue that comparing *ω*_A_ to 0 is not a robust test for adaptation. Instead, *ω*_A_ for a particular genomic region of interest should be compared to other genomic regions for which the nearly-neutral evolution is not rejected, and the difference Δ*ω*_A_ should be compared to 0, as done in this study. More generally, our empirical analysis emphasizes the limitations of, and the difficulties raised by, the model-based population-genetic approaches. In this respect, further exploring the congruence (or lack thereof) between phylogenetic and population-genetic approaches will represent a useful asset to clarify those delicate problems, given that similar benefits are also expected on the side of phylogenetic approaches, which are far from immune from methodological limitations.

More broadly on a theoretical level, this work leverages a specific overlap between phylogenetic and population genetics, namely that the rate of adaptation 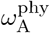 in phylogenetic codon models and *ω*_A_ in the MK test should theoretically be directly comparable. Based on this theoretical relationship, our study is paving the way for studies and methods augmenting molecular polymorphism data within species with information about divergence data between species[40], and by assessing empirically the relationship between phylogenetic and population genetics[41]. In this light, mutation-selection models at the phylogenetic scale can play a dual role: pinpointing genes and sites under adaptation 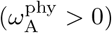, and also seeking the genomic region for which the nearly-neutral theory is not rejected 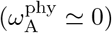.

## Methods

### Phylogenetic dataset

Protein-coding DNA sequences alignments in placental mammals and their corresponding gene trees were extracted from the [OrthoMaM](https://orthomam.mbb.cnrs.fr/) database, containing 116 mammalian reference sequences in v10c[42–44]. Genes located on the X, Y and mitochondrial chromosome were discarded from the analysis, since the number of polymorphism, necessary in population-based method, is expected to be different on these sequences. Additionally, sequences from the species for which polymorphism are available, as well as their sister species have been discarded from the analysis to ensure independence between the data used in the phylogenetic and population-genetic method. Altogether, we analyzed 14,509 protein-coding DNA sequences alignment containing at most 87 reference sequences of placental mammals.

### Adaptation in phylogeny-based method

Classical codon models estimates a parameter *ω* = *d*_*N*_ */d*_*S*_, namely the ratio of the non-synonymous over the synonymous substitution rates[4, 5]. In the so-called site models, *ω* is allowed to vary across sites[11, 45]. In *Bayescode*, site-specific *ω*^(*i*)^ (fig. 1B, y-axis) are independent identically distributed from a gamma distribution[46]. In a second step, the average over sites is calculated, giving estimates of *ω* for each protein-coding sequence (fig. 1A, y-axis).

In contrast, mutation-selection models assume that the protein-coding sequence is at mutation-selection balance under a fixed fitness landscape, which is itself characterized by a fitness vector over the 20 amino acid at each site[13–15]. Mathematically, the rate of non-synonymous substitution from codon *a* to codon *b* 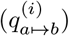at site *i* of the sequence is equal to the rate of mutation from the underlying DNA change (*μ*_*a b*_) multiplied by the scaled probability of fixation of the mutation 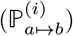 Crucially, the probability of fixation depends on the difference of scaled fitness between the amino acid encoded by the mutated codon 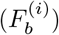 and the fitness of the amino acid encoded by the original codon 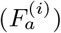of site *i*[47, 48]. Altogether, the rate of substitution from codon *a* to *b* at a given site *i* is:

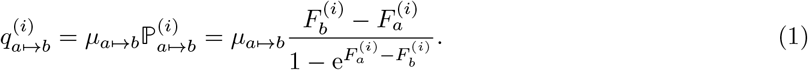

Fitting the mutation-selection model on a sequence alignment leads to an estimation of the mutation rate matrix (***μ***) as well as the 20 amino acid fitness landscape (***F*** ^**(*i*)**^) at each site *i*. From these parameters, one can compute 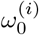 (fig. 1B, x-axis), the site-specific rate of non-synonymous over synonymous substitution at the mutation-selection balance:

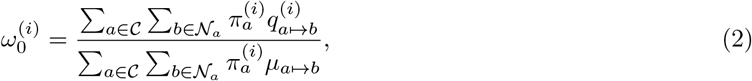

where 𝒞 is the set all the possible codons (61 by discarding stop codons), 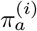is the equilibrium frequency of codon *a* at site *i*, and 𝒩_*a*_ is the set of codons that are non-synonymous to *a*[16, 17]. The equilibrium frequency of codon *a* at site *i* is the product of the nucleotide frequencies at its three positions and the scaled Wrightian fitness of the amino acid 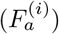:

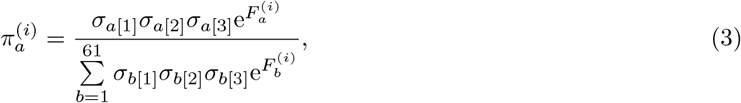

where *σ*_*a*[*j*]_ *∈ {A, T, C, G}* is the equilibrium frequency (given by the mutational matrix) of the nucleotide at position *j ∈ {*1, 2, 3*}* of codon *a*. In a second step, the average over sites is calculated, giving estimates of *ω*_0_ for each protein-coding sequences (fig. 1A, x-axis). Under the assumption that the protein is under a nearly-neutral regime, the calculated *ω*_0_ (mutation-selection model) and the estimated *ω* (site model) should be the same[16].

We ran the Bayesian software *BayesCode* (https://github.com/ThibaultLatrille/bayescode) on each protein-coding DNA alignment[49]. Each Monte-Carlo Markov-Chain (MCMC) is run during 2,000 points, with a burn-in of 1,000 points, to obtain the posterior mean of *ω* and *ω*_0_ across the MCMC, as well as the 95% posterior credibility interval for genes and sites. Genes and sites classified under an adaptive regime (in red) are rejecting the nearly-neutral assumption such that the lower bound for the credible interval of *ω* (*α* = 0.05) is above the upper bound of the credible interval of *ω*_0_ (*α* = 0.05), meaning that the value of their *ω* is higher than that of their *ω*_0_. Because this is a unilateral test (*ω > ω*_0_) and the two credible interval are independent, the risk is (*α/*2)^2^ = 0.025^2^ = 0.000625 for each test. Empirically, the nearly-neutral assumption appears to be rejected for 822 out 14,509 genes, while 0.000625 *×* 14,509 *≃* 9 genes are expected due to the multiple testing, suggesting a 9*/*822 *≃* 1% rate of false positive at the gene level. At the site level, the nearly-neutral assumption appears to be rejected for 104,129 out of 8,895,374 sites, while 0.000625 *×* 8,895,374 *≃* 5,560 are expected due to the multiple testing, suggesting a 5,560*/*104,129 *≃* 5% rate of false positive at the site level. Genes and sites are classified under a nearly-neutral regime (in green) if the average *ω* is within the credible interval of the *ω*_0_, and respectively the average *ω*_0_ is also within the credible interval of *ω*, meaning *ω* = *ω*_0_. Additionally, the set of sites detected exclusively by mutation-selection codon models have a mean *ω <* 1. Genes and sites that do not fall in any of these categories are considered unclassified.

### Polymorphism dataset

Each SNP (chromosome, position, strand) in the focal species was matched to its relative position (chromosome, position, strand) in the protein-coding DNA alignment by first converting the genomic positions to relative position in the coding sequence (CDS) using gene annotation files (GTF format) downloaded from Ensembl (ensembl.org). We then verified that the SNP downloaded from Ensembl were matching the reference in the CDS (FASTA format). Second, the relative position in the CDS was converted to position in the multiple sequence alignment (containing gaps) from OrthoMaM database[42–44] by doing a global pairwise alignment, using the Biopython function pairwise2, between the CDS fasta and the sequence found in the alignment. This conversion from genomic position to position in the alignment is only possible if the assembly used for SNP calling is the same as the one used in the alignment, the GTF annotations and the FASTA sequences.

We retrieved the genetic variants representing the population level polymorphism from the following species and respective available datasets: *Equus caballus* (EquCab2 assembly in the EVA study PRJEB9799[50]), *Canis familiars* (CanFam3.1 assembly in the EVA study PRJEB24066[51]), *Bos taurus* (UMD3.1 assembly in the NextGen project), *Ovis aries* (Oar v3.1 assembly in the NextGen project), *Capra Hircus* (CHIR1 assembly in the NextGen project converted to ARS1 assembly with dbSNP identifiers[52]), *Chlorocebus sabaeus* (ChlSab1.1 assembly in the EVA project PRJEB22989[53]), *Homo sapiens* (GRCh38 assembly from the 1000-genome project[54, 55]).

Variants not inside genes are discarded at the beginning of the analysis. Insertions and deletions are not analyzed, and only Single Nucleotide Polymorphisms (SNPs) with only one mutant allele are considered. Stop codon mutants are also discarded. For populations containing more than 8 sampled individuals, the site-frequency spectrum (SFS) is subsampled down to 16 chromosomes (8 diploid individuals) without replacement (hyper-geometric distribution) to alleviate the effect of different sampling depth in the 29 populations. Moreover, subsampling mitigate the impact of moderately deleterious mutations segregating at low frequency on *π*_*N*_ */π*_*S*_, since they are more likely to be discarded than polymorphism segregating at higher frequency. The Snakemake pipeline for integrating polymorphism and divergence data uses custom scripts written in python 3.9.

### Rate of adaption in population-based method

The genes and sites classified as under adaptation are concatenated. For each population *π*_*N*_ */π*_*S*_ is computed as the sum of non-synonymous over synonymous polymorphism on the concatenated SFS. *d*_*N*_ */d*_*S*_ is computed on the concatenated pairwise alignment between focal and sister species extracted from OrthoMaM, the *d*_*N*_ */d*_*S*_ count is performed by *yn00*. We considered *Ceratotherium simum simum* as *Equus caballus* sister species; *Ursus maritimus* as *Canis familiars* sister species; *Bison bison bison* as *Bos taurus* sister species; *Pantholops hodgsonii* as *Ovis aries* sister species; *Pantholops hodgsonii* as *Capra Hircus* sister species; *Macaca mulatta* as *Chlorocebus sabaeus* sister species and finally we considered *Pan troglodytes* as *Homo sapiens* sister species. Altogether, *ω*_A_ = *d*_*N*_ */d*_*S*_ *− π*_*N*_ */π*_*S*_ is thus computed for each population on genes and sites classified as under adaptation. The result is compared to the empirical null distribution of *ω*_A_, obtained by randomly sampling (1,000 sampling replicates) a subset of genes/sites classified as nearly-neutral.

Other methods to compute *ω*_A_ such as polyDFE[20] are also used (eq. 3-20 in supplementary materials), which relies on the synonymous and non-synonymous unfolded site-frequency spectra (SFS) to estimate the distribution of fitness effects of mutations (DFE), and the rate of adaptation. In polyDFE, GammaExpo models the fitness effect of weakly deleterious non-synonymous mutations as distributed according to a negative Gamma and the fitness effect of weakly advantageous mutations are distributed exponentially. This method is an extension of the methods introduced by Eyre-Walker and collaborators[9, 56]. Unfolded SFSs are obtained by polarizing SNPs using the 3 closest outgroups found in the OrthoMam alignment with est-usfs v2.04[57].

## 1 Data availability

The data underlying this article are available at 10.5281/zenodo.7107234. Scripts and instructions necessary to reproduce the empirical experiments on the original dataset or with user-specified datasets is available at https://github.com/ThibaultLatrille/AdaptaPop.

## 2 Acknowledgements

We gratefully also acknowledge the help of Nicolas Galtier and Julien Joseph for their advice and review concerning this manuscript. This work was performed using the computing facilities of the CC LBBE/PRABI. This study makes use of data generated by the NextGen Consortium. The European Union’s Seventh Framework Programme (FP7/2010-2014) provided funding for the project under grant agreement no 244356 - “NextGen”. Funding: Université de Lausanne. Agence Nationale de la Recherche, Grant ANR-15-CE12-0010-01 / DASIRE. Agence Nationale de la Recherche, Grant ANR-19-CE12-0019 / HotRec.

## 3 Author information

TL, NR and NL designed the study. TL gathered and formatted the data and conducted the analyses with BayesCode using scripts in Python and pipeline in Snakemake. TL, NR and NL contributed to the writing of the manuscript.

## Notes

https://github.com/ThibaultLatrille/AdaptaPop

https://zenodo.org/record/7107234

### Competing Interest Statement

The authors have declared no competing interest.

