## Supplementary materials for "Genes and sites under adaptation at the phylogenetic scale also exhibit adaptation at the population-genetic scale"

#### Contents

**1 Running Bayescode** **2**

1.1 Site-specific $\omega$-based codon models. 2

1.2 Site-specific mutation-selection codon models 2

**2 Gene ontology enrichment at gene and site level** **3**

2.1 Gene-specific mutation-selection model 3

2.2 Site-specific $\boldsymbol{\omega}$-based model 4

2.3 Site-specific mutation-selection model 5

2.4 Exclusive to site-specific mutation-selection model 7

**3 Graphical abstract for the pipeline.** **9**

**4 Rate of adaptation enrichment while controlling for $\boldsymbol{\omega}$** **10**

4.1 Mutation-selection codon model at gene level ($\boldsymbol{\alpha = 0.05}$) 11

4.2 Mutation-selection codon model at site level ($\boldsymbol{\alpha = 0.05}$) 13

**5 Rate of adaptation enrichment with $\boldsymbol{\alpha = 0.005}$** **15**

5.1 Scatterplot with $\boldsymbol{\alpha = 0.005}$ 15

5.2 Mutation-selection codon model at gene level ($\boldsymbol{\alpha = 0.005}$) 16

5.3 Mutation-selection codon model at site level ($\boldsymbol{\alpha = 0.005}$) 17

**6 Rate of adaptation enrichment with polyDFE** **18**

6.1 PolyDFE model C - including divergence data 18

6.2 PolyDFE model C - polymorphism data alone with $\beta > 0$ 21

6.3 PolyDFE model C - polymorphism data alone with $\beta > 5$ 23

6.4 PolyDFE model D - including divergence 25

### 1 Running Bayescode

#### 1.1 Site-specific $\omega$-based codon models.

The 61-by-61 codon substitution matrix ($\boldsymbol{q^{(i)}}$) at site $i$ is defined entirely by the mutation matrix ($\boldsymbol{\mu}$), $\omega^{(i)}$ and the genetic code:

$$\begin{cases} q^{(i)}_{a \mapsto b} & = 0 \text{ if codons } a \text{ and } b \text{ are more than one mutation away,} \\ q^{(i)}_{a \mapsto b} & = \mu_{a \mapsto b} \text{ if codons } a \text{ and } b \text{ are synonymous,} \\ q^{(i)}_{a \mapsto b} & = \omega^{(i)} \mu_{a \mapsto b} \text{ if codons } a \text{ and } b \text{ are non-synonymous.} \end{cases} \tag{1}$$

By definition of the instantaneous rate matrix, the sum of the entries in each row of the codon substitution rate matrix $\boldsymbol{q}$ is equal to 0, giving the diagonal entries:

$$q^{(i)}_{a \mapsto b} = - \sum_{b \neq a, b=1}^{61} q^{(i)}_{a \mapsto b}. \tag{2}$$

In BayesCode (https://github.com/ThibaultLatrille/bayescode), $\omega$-based site-specific codon models are obtained by running *mutselomega* with the options:

```
mutselomega --omegashift 0.0 --freeomega --omegancat 30 --flatfitness -a my_alignment.phy -t my_tree.newick -u 2000 my_genename
```

The mean value of $\omega$ per site is then obtained by running *readmutselomega* with the options:

```
readmutselomega --every 1 --until 2000 --burnin 1000 -c 0.025 my_genename
```

#### 1.2 Site-specific mutation-selection codon models

In BayesCode (https://github.com/ThibaultLatrille/bayescode), mutation-selection codon models are obtained by running *mutselomega* for 2000 points of MCMC with the options:

```
mutselomega ---omegashift 0.0 --ncat 30 -a my_alignment.phy -t my_tree.newick -u 2000 my_genename
```

The collection of site-specific fitness profiles ($\boldsymbol{F^{(i)}}, \forall i$) are then obtained by running *readmutselomega*, reading 1000 points of MCMC (first 1000 are considered as burn-in) with the options:

```
readmutselomega --every 1 --until 2000 --burnin 1000 --ss my_genename
```

The gene-specific mutation matrix ($\boldsymbol{\mu}$) is also obtained by running *readmutselomega*, reading 1000 points of MCMC (first 1000 are considered as burn-in) with the options:

```
readmutselomega --every 1 --until 2000 --burnin 1000 --nuc my_genename
```

#### 2 Gene ontology enrichment at gene and site level

##### 2.1 Gene-specific mutation-selection model

Genes are classified under an adaptive regime if the lower bound for the posterior credibility interval of gene-specific $\omega$ ($\alpha = 0.05$) is above the upper bound of the posterior credibility interval of gene-specific $\omega_0$. Because this is a unilateral test ($\omega > \omega_0$) and the two posterior credibility interval are independent, the risk is $(\alpha/2)^2 = 0.025^2 = 6.25 \times 10^{-4}$ for each gene. Genes are classified as control if they are not in the adaptive group. For each ontology (775 ontologies), a 2x2 contingency tables is built by counting the number of genes based on their evolutionary regime (adaptive regime or control group) and their ontology (whether they have this specific ontology or not). Fisher's exact tests are then performed for these 2x2 contingency tables. $p_{\text{v}}^{\text{adj}}$ are corrected for multiple comparison (Holm–Bonferroni correction).

Table S1: Ontology enrichment with gene-specific mutation-selection model

| Gene ontology | $n_{\text{Observed}}$ | $n_{\text{Expected}}$ | Odds ratio | $p_{\text{v}}$ | $p_{\text{v}}^{\text{adj}}$ (* if $< 0.05$) |
|---|---|---|---|---|---|
| immune system process | 48 | 5.625 | 8.533 | $9.8 \times 10^{-24}$ | $\mathbf{7.6 \times 10^{-21}}$* |
| extracellular space | 82 | 19.8 | 4.151 | $6.2 \times 10^{-21}$ | $\mathbf{4.8 \times 10^{-18}}$* |
| innate immune response | 41 | 5.245 | 7.817 | $1.6 \times 10^{-19}$ | $\mathbf{1.2 \times 10^{-16}}$* |
| extracellular region | 99 | 30.5 | 3.251 | $5 \times 10^{-18}$ | $\mathbf{3.9 \times 10^{-15}}$* |
| regulation of complement activation | 14 | 0.359 | 39.0 | $1.2 \times 10^{-13}$ | $\mathbf{9.4 \times 10^{-11}}$* |
| extracellular exosome | 100 | 38.2 | 2.618 | $3.6 \times 10^{-13}$ | $\mathbf{2.7 \times 10^{-10}}$* |
| complement activation | 12 | 0.180 | 66.6 | $5.3 \times 10^{-13}$ | $\mathbf{4 \times 10^{-10}}$* |
| immune response | 28 | 4.054 | 6.907 | $6.5 \times 10^{-13}$ | $\mathbf{5 \times 10^{-10}}$* |
| blood microparticle | 17 | 1.247 | 13.6 | $7.1 \times 10^{-12}$ | $\mathbf{5.5 \times 10^{-9}}$* |
| serine-type endopeptidase activity | 20 | 2.243 | 8.917 | $3.1 \times 10^{-11}$ | $\mathbf{2.4 \times 10^{-8}}$* |
| inflammatory response | 27 | 5.129 | 5.264 | $2.5 \times 10^{-10}$ | $\mathbf{1.9 \times 10^{-7}}$* |
| integral component of membrane | 149 | 78.4 | 1.901 | $1.4 \times 10^{-8}$ | $\mathbf{1.1 \times 10^{-5}}$* |
| defense response to virus | 15 | 1.795 | 8.355 | $1.7 \times 10^{-8}$ | $\mathbf{1.3 \times 10^{-5}}$* |
| complement activation | 8 | 0.182 | 43.9 | $1.8 \times 10^{-8}$ | $\mathbf{1.4 \times 10^{-5}}$* |
| integral component of plasma membrane | 58 | 23.0 | 2.516 | $2 \times 10^{-8}$ | $\mathbf{1.5 \times 10^{-5}}$* |
| leukocyte migration | 16 | 2.210 | 7.239 | $2.8 \times 10^{-8}$ | $\mathbf{2.1 \times 10^{-5}}$* |
| neutrophil degranulation | 28 | 7.432 | 3.767 | $6 \times 10^{-8}$ | $\mathbf{4.5 \times 10^{-5}}$* |
| external side of plasma membrane | 18 | 3.156 | 5.703 | $7.6 \times 10^{-8}$ | $\mathbf{5.7 \times 10^{-5}}$* |
| plasma membrane | 131 | 70.7 | 1.852 | $1.2 \times 10^{-7}$ | $\mathbf{9.1 \times 10^{-5}}$* |
| lipid metabolic process | 31 | 9.452 | 3.280 | $1.9 \times 10^{-7}$ | **0.00014*** |
| proteolysis | 34 | 11.2 | 3.047 | $2.4 \times 10^{-7}$ | **0.00018*** |
| platelet degranulation | 14 | 1.982 | 7.064 | $2.5 \times 10^{-7}$ | **0.00019*** |
| cell surface | 29 | 9.210 | 3.149 | $9.1 \times 10^{-7}$ | **0.00069*** |
| extracellular vesicle | 10 | 0.969 | 10.3 | $9.5 \times 10^{-7}$ | **0.00072*** |
| serine-type peptidase activity | 13 | 1.927 | 6.746 | $1 \times 10^{-6}$ | **0.00076*** |
| toll-like receptor signaling pathway | 9 | 0.729 | 12.4 | $1.1 \times 10^{-6}$ | **0.00086*** |
| chemotaxis | 13 | 1.988 | 6.540 | $1.3 \times 10^{-6}$ | **0.001*** |
| defense response to bacterium | 12 | 1.690 | 7.100 | $1.7 \times 10^{-6}$ | **0.001*** |
| adaptive immune response | 13 | 2.048 | 6.347 | $1.7 \times 10^{-6}$ | **0.001*** |
| cilium movement | 7 | 0.366 | 19.1 | $2.8 \times 10^{-6}$ | **0.002*** |
| apical plasma membrane | 21 | 5.830 | 3.602 | $4.2 \times 10^{-6}$ | **0.003*** |
| chemokine-mediated signaling pathway | 7 | 0.488 | 14.3 | $9.5 \times 10^{-6}$ | **0.007*** |
| cell surface receptor signaling pathway | 18 | 4.787 | 3.760 | $1.2 \times 10^{-5}$ | **0.009*** |
| cytolysis | 5 | 0.123 | 40.8 | $1.3 \times 10^{-5}$ | **0.009*** |
| antimicrobial humoral response | 5 | 0.123 | 40.8 | $1.3 \times 10^{-5}$ | **0.009*** |
| positive regulation of heterotypic cell-cell adhesion | 5 | 0.123 | 40.8 | $1.3 \times 10^{-5}$ | **0.009*** |
| hemostasis | 9 | 1.094 | 8.225 | $1.3 \times 10^{-5}$ | **0.010*** |
| cellular protein metabolic process | 14 | 3.134 | 4.468 | $2 \times 10^{-5}$ | **0.015*** |
| positive regulation of ERK1 and ERK2 cascade | 14 | 3.194 | 4.383 | $2.5 \times 10^{-5}$ | **0.018*** |
| platelet alpha granule | 5 | 0.184 | 27.2 | $3.2 \times 10^{-5}$ | **0.024*** |
| dynein light chain binding | 5 | 0.184 | 27.2 | $3.2 \times 10^{-5}$ | **0.024*** |
| lysosome | 21 | 7.167 | 2.930 | $6.1 \times 10^{-5}$ | **0.045*** |
| negative regulation of viral genome replication | 5 | 0.245 | 20.4 | $6.9 \times 10^{-5}$ | 0.051 |
| regulation of immune response | 8 | 1.097 | 7.289 | $7.9 \times 10^{-5}$ | 0.057 |
| receptor activity | 15 | 4.158 | 3.607 | $8.8 \times 10^{-5}$ | 0.064 |
| peptidase activity | 25 | 9.815 | 2.547 | 0.0001 | 0.074 |
| receptor binding | 21 | 7.473 | 2.810 | 0.0001 | 0.075 |

Continued on next page

| Gene ontology | $n_{\text{Observed}}$ | $n_{\text{Expected}}$ | Odds ratio | $p_{\text{v}}$ | $p_{\text{v}}^{\text{adj}}$ (* if < 0.05) |
|---|---|---|---|---|---|
| membrane | 185 | 122.7 | 1.508 | 0.00013 | 0.095 |
| fibrinolysis | 5 | 0.307 | 16.3 | 0.00013 | 0.096 |
| negative regulation of blood coagulation | 4 | 0.123 | 32.5 | 0.00016 | 0.115 |
| neutrophil chemotaxis | 6 | 0.612 | 9.797 | 0.00019 | 0.136 |
| defense response to Gram-negative bacterium | 6 | 0.612 | 9.797 | 0.00019 | 0.136 |
| receptor-mediated endocytosis | 11 | 2.549 | 4.315 | 0.0002 | 0.141 |
| hydrolase activity | 54 | 29.9 | 1.804 | 0.00021 | 0.151 |
| acrosomal membrane | 5 | 0.368 | 13.6 | 0.00023 | 0.166 |
| response to virus | 8 | 1.342 | 5.960 | 0.00024 | 0.172 |
| antioxidant activity | 4 | 0.185 | 21.7 | 0.00035 | 0.253 |
| natural killer cell activation | 4 | 0.185 | 21.7 | 0.00035 | 0.253 |
| plasminogen activation | 4 | 0.185 | 21.7 | 0.00035 | 0.253 |
| positive regulation of angiogenesis | 10 | 2.312 | 4.326 | 0.00037 | 0.264 |
| positive regulation of phagocytosis | 5 | 0.430 | 11.6 | 0.00038 | 0.269 |
| negative regulation of endopeptidase activity | 8 | 1.465 | 5.461 | 0.00039 | 0.277 |
| blood coagulation | 12 | 3.336 | 3.597 | 0.00043 | 0.308 |
| organelle membrane | 8 | 1.526 | 5.242 | 0.00049 | 0.346 |
| cellular response to interleukin-1 | 6 | 0.797 | 7.532 | 0.00055 | 0.388 |
| fatty acid metabolic process | 11 | 3.039 | 3.620 | 0.0007 | 0.499 |
| platelet alpha granule lumen | 7 | 1.223 | 5.721 | 0.00071 | 0.505 |
| defense response | 6 | 0.858 | 6.993 | 0.00074 | 0.526 |
| specific granule membrane | 8 | 1.772 | 4.516 | 0.001 | 0.778 |
| bile acid biosynthetic process | 4 | 0.308 | 13.0 | 0.001 | 0.813 |
| phospholipase A2 activity | 4 | 0.308 | 13.0 | 0.001 | 0.813 |
| cholesterol efflux | 4 | 0.308 | 13.0 | 0.001 | 0.813 |
| apoptotic cell clearance | 4 | 0.308 | 13.0 | 0.001 | 0.813 |
| arachidonic acid secretion | 4 | 0.308 | 13.0 | 0.001 | 0.813 |

The genes detected under adaptation by mutation-selection codon models are enriched primarily with ontologies related to immune system processes (innate immune response, immune response, inflammatory response, defense response to virus, etc) and ontologies related to the external membrane (extracellular region, extracellular exosome, etc).

#### 2.2 Site-specific $\omega$-based model

For each gene, we computed the proportion of sites classified under an adaptive regime such that the lower bound for the posterior credibility interval of site-specific $\omega$ ($\alpha = 0.05$) is above 1. Because this is a unilateral test ($\omega > 1$) the risk is $\alpha/2 = 0.05/2 = 0.025$ for each site. For each ontology, the proportion of sites under adaptation is compared between the set of genes sharing this given ontology and the rest of the genes with Mann-Whitney U test. $p_{\text{v}}^{\text{adj}}$ are corrected for multiple comparison (Holm–Bonferroni correction).

Table S2: Ontology enrichment with site-specific $\omega$-based model

| Gene ontology | Mann-Whitney U | $p_{\text{v}}$ | $p_{\text{v}}^{\text{adj}}$ (* if < 0.05) |
|---|---|---|---|
| extracellular region | $2.1 \times 10^{6}$ | $1 \times 10^{-17}$ | $\mathbf{5.1 \times 10^{-15}}$* |
| extracellular space | $1.5 \times 10^{6}$ | $3.9 \times 10^{-15}$ | $\mathbf{2 \times 10^{-12}}$* |
| immune system process | $5.6 \times 10^{5}$ | $7.5 \times 10^{-15}$ | $\mathbf{3.7 \times 10^{-12}}$* |
| immune response | $3.7 \times 10^{5}$ | $2 \times 10^{-12}$ | $\mathbf{9.8 \times 10^{-10}}$* |
| external side of plasma membrane | $2.8 \times 10^{5}$ | $8.1 \times 10^{-11}$ | $\mathbf{4 \times 10^{-8}}$* |
| innate immune response | $4.8 \times 10^{5}$ | $6.9 \times 10^{-10}$ | $\mathbf{3.4 \times 10^{-7}}$* |
| blood microparticle | $1.6 \times 10^{5}$ | $2.6 \times 10^{-9}$ | $\mathbf{1.3 \times 10^{-6}}$* |
| inflammatory response | $4.1 \times 10^{5}$ | $2.1 \times 10^{-8}$ | $\mathbf{1 \times 10^{-5}}$* |
| DNA repair | $4.4 \times 10^{5}$ | $1.6 \times 10^{-6}$ | **0.0008*** |
| lipid metabolic process | $6.4 \times 10^{5}$ | $1.8 \times 10^{-6}$ | **0.00086*** |
| serine-type endopeptidase activity | $2.2 \times 10^{5}$ | $1.9 \times 10^{-6}$ | **0.0009*** |
| receptor activity | $3 \times 10^{5}$ | $3.3 \times 10^{-6}$ | **0.002*** |
| cytokine activity | $2.4 \times 10^{5}$ | $3.7 \times 10^{-6}$ | **0.002*** |
| apical plasma membrane | $4.1 \times 10^{5}$ | $4.7 \times 10^{-6}$ | **0.002*** |

Continued on next page

| Gene ontology | Mann-Whitney U | $p_{\text{v}}$ | $p_{\text{v}}^{\text{adj}}$ (* if < 0.05) |
|---|---|---|---|
| cell surface receptor signaling pathway | $3.4 \times 10^{5}$ | $5.9 \times 10^{-6}$ | **0.003*** |
| integral component of plasma membrane | $1.4 \times 10^{6}$ | $1.1 \times 10^{-5}$ | **0.005*** |
| fatty acid metabolic process | $2.2 \times 10^{5}$ | $1.1 \times 10^{-5}$ | **0.005*** |
| cell surface | $6.1 \times 10^{5}$ | $1.2 \times 10^{-5}$ | **0.006*** |
| defense response to bacterium | $1.5 \times 10^{5}$ | $1.5 \times 10^{-5}$ | **0.007*** |
| centriole | $1.7 \times 10^{5}$ | $1.5 \times 10^{-5}$ | **0.007*** |
| leukocyte migration | $2 \times 10^{5}$ | $1.5 \times 10^{-5}$ | **0.007*** |
| proteolysis | $7.2 \times 10^{5}$ | $2 \times 10^{-5}$ | **0.009*** |
| cell-matrix adhesion | $1.5 \times 10^{5}$ | $2 \times 10^{-5}$ | **0.009*** |
| defense response to virus | $1.7 \times 10^{5}$ | $2.1 \times 10^{-5}$ | **0.010*** |
| cellular response to DNA damage stimulus | $5.6 \times 10^{5}$ | $3.7 \times 10^{-5}$ | **0.017*** |
| peroxisome | $1.7 \times 10^{5}$ | $3.7 \times 10^{-5}$ | **0.018*** |
| extracellular exosome | $2.3 \times 10^{6}$ | $3.9 \times 10^{-5}$ | **0.018*** |
| viral entry into host cell | $1.2 \times 10^{5}$ | $6.1 \times 10^{-5}$ | **0.028*** |
| lipid catabolic process | $1.6 \times 10^{5}$ | $7.5 \times 10^{-5}$ | **0.035*** |
| cellular protein metabolic process | $2.3 \times 10^{5}$ | $9.6 \times 10^{-5}$ | **0.045*** |
| integral component of membrane | $3.8 \times 10^{6}$ | 0.00015 | 0.068 |
| metallopeptidase activity | $2.6 \times 10^{5}$ | 0.00017 | 0.081 |
| serine-type peptidase activity | $1.6 \times 10^{5}$ | 0.00019 | 0.088 |
| platelet degranulation | $1.7 \times 10^{5}$ | 0.00024 | 0.110 |
| cilium | $3.3 \times 10^{5}$ | 0.00026 | 0.118 |
| peptidase activity | $6.1 \times 10^{5}$ | 0.00026 | 0.121 |
| neutrophil degranulation | $5 \times 10^{5}$ | 0.00029 | 0.135 |
| cell adhesion | $6.2 \times 10^{5}$ | 0.00033 | 0.152 |
| ciliary basal body-plasma membrane docking | $1.4 \times 10^{5}$ | 0.00046 | 0.209 |
| DNA replication | $2 \times 10^{5}$ | 0.00053 | 0.243 |
| meiotic cell cycle | $1.7 \times 10^{5}$ | 0.00056 | 0.254 |
| collagen trimer | $1.3 \times 10^{5}$ | 0.00064 | 0.291 |
| receptor-mediated endocytosis | $1.9 \times 10^{5}$ | 0.00069 | 0.314 |
| carbohydrate binding | $1.9 \times 10^{5}$ | 0.0007 | 0.317 |
| regulation of G2/M transition of mitotic cell cycle | $1.2 \times 10^{5}$ | 0.001 | 0.516 |
| steroid metabolic process | $1.4 \times 10^{5}$ | 0.002 | 0.694 |
| receptor binding | $4.7 \times 10^{5}$ | 0.002 | 0.814 |
| positive regulation of ERK1 and ERK2 cascade | $2.3 \times 10^{5}$ | 0.002 | 0.884 |

The sites detected under adaptation by site-specific $\omega$-based codon models are enriched primarily with ontologies related to immune system processes (innate immune response, immune response, inflammatory response, defense response to virus, etc) and ontologies related to the external membrane (extracellular region, extracellular exosome, etc).

#### 2.3 Site-specific mutation-selection model

For each gene, we computed the proportion of sites classified under an adaptive regime such that the lower bound for the posterior credibility interval of site-specific $\omega$ ($\alpha = 0.05$) is above the upper bound of the posterior credibility interval of site-specific $\omega_0$ ($\alpha = 0.05$). Because this is a unilateral test ($\omega > \omega_0$) and the two posterior credibility interval are independent, the risk is $(\alpha/2)^2 = 0.025^2 = 6.25 \times 10^{-4}$ for each site. For each ontology, the proportion of sites under adaptation is compared between the set of genes sharing this given ontology and the rest of the genes with Mann-Whitney U test. $p_{\text{v}}^{\text{adj}}$ are corrected for multiple comparison (Holm–Bonferroni correction).

Table S3: Ontology enrichment with site-specific mutation-selection model

| Gene ontology | Mann-Whitney U | $p_{\text{v}}$ | $p_{\text{v}}^{\text{adj}}$ (* if < 0.05) |
|---|---|---|---|
| extracellular space | $1.6 \times 10^{6}$ | $8.4 \times 10^{-22}$ | $\mathbf{4.1 \times 10^{-19}}$* |
| extracellular region | $2.2 \times 10^{6}$ | $2.8 \times 10^{-21}$ | $\mathbf{1.4 \times 10^{-18}}$* |
| lipid metabolic process | $7.7 \times 10^{5}$ | $3.4 \times 10^{-21}$ | $\mathbf{1.7 \times 10^{-18}}$* |
| oxidation-reduction process | $8.8 \times 10^{5}$ | $1.6 \times 10^{-19}$ | $\mathbf{8 \times 10^{-17}}$* |
| integral component of membrane | $4.1 \times 10^{6}$ | $1.8 \times 10^{-18}$ | $\mathbf{8.6 \times 10^{-16}}$* |
| oxidoreductase activity | $7.8 \times 10^{5}$ | $3.2 \times 10^{-17}$ | $\mathbf{1.6 \times 10^{-14}}$* |

Continued on next page

| Gene ontology | Mann-Whitney U | $p_{\text{v}}$ | $p_{\text{v}}^{\text{adj}}$ (* if < 0.05) |
|---|---|---|---|
| extracellular exosome | $2.5 \times 10^{6}$ | $1.6 \times 10^{-15}$ | $\mathbf{7.8 \times 10^{-13}}$* |
| immune system process | $5.9 \times 10^{5}$ | $9.1 \times 10^{-14}$ | $\mathbf{4.5 \times 10^{-11}}$* |
| immune response | $4.1 \times 10^{5}$ | $2.1 \times 10^{-13}$ | $\mathbf{1 \times 10^{-10}}$* |
| integral component of plasma membrane | $1.5 \times 10^{6}$ | $5.4 \times 10^{-13}$ | $\mathbf{2.6 \times 10^{-10}}$* |
| inflammatory response | $4.6 \times 10^{5}$ | $3.3 \times 10^{-12}$ | $\mathbf{1.6 \times 10^{-9}}$* |
| fatty acid metabolic process | $2.7 \times 10^{5}$ | $1.3 \times 10^{-11}$ | $\mathbf{6.2 \times 10^{-9}}$* |
| innate immune response | $5.2 \times 10^{5}$ | $1.1 \times 10^{-10}$ | $\mathbf{5.3 \times 10^{-8}}$* |
| mitochondrion | $1.6 \times 10^{6}$ | $1.6 \times 10^{-10}$ | $\mathbf{7.5 \times 10^{-8}}$* |
| serine-type endopeptidase activity | $2.5 \times 10^{5}$ | $1.6 \times 10^{-10}$ | $\mathbf{7.8 \times 10^{-8}}$* |
| metabolic process | $5.9 \times 10^{5}$ | $1.7 \times 10^{-10}$ | $\mathbf{8.1 \times 10^{-8}}$* |
| mitochondrial inner membrane | $5 \times 10^{5}$ | $4.8 \times 10^{-10}$ | $\mathbf{2.3 \times 10^{-7}}$* |
| blood microparticle | $1.8 \times 10^{5}$ | $5 \times 10^{-10}$ | $\mathbf{2.4 \times 10^{-7}}$* |
| external side of plasma membrane | $2.9 \times 10^{5}$ | $4.3 \times 10^{-9}$ | $\mathbf{2.1 \times 10^{-6}}$* |
| defense response to bacterium | $1.8 \times 10^{5}$ | $3.9 \times 10^{-8}$ | $\mathbf{1.8 \times 10^{-5}}$* |
| catalytic activity | $6.5 \times 10^{5}$ | $7.5 \times 10^{-8}$ | $\mathbf{3.6 \times 10^{-5}}$* |
| symporter activity | $1.8 \times 10^{5}$ | $1.1 \times 10^{-7}$ | $\mathbf{5.4 \times 10^{-5}}$* |
| membrane | $4.7 \times 10^{6}$ | $1.2 \times 10^{-7}$ | $\mathbf{5.5 \times 10^{-5}}$* |
| peroxisome | $1.9 \times 10^{5}$ | $1.2 \times 10^{-7}$ | $\mathbf{5.6 \times 10^{-5}}$* |
| cell surface | $6.5 \times 10^{5}$ | $1.2 \times 10^{-7}$ | $\mathbf{5.7 \times 10^{-5}}$* |
| apical plasma membrane | $4.4 \times 10^{5}$ | $1.4 \times 10^{-7}$ | $\mathbf{6.6 \times 10^{-5}}$* |
| serine-type peptidase activity | $1.9 \times 10^{5}$ | $4.4 \times 10^{-7}$ | **0.00021*** |
| hydrolase activity | $1.7 \times 10^{6}$ | $5.7 \times 10^{-7}$ | **0.00027*** |
| steroid metabolic process | $1.7 \times 10^{5}$ | $6.6 \times 10^{-7}$ | **0.00031*** |
| neutrophil degranulation | $5.5 \times 10^{5}$ | $7.7 \times 10^{-7}$ | **0.00036*** |
| flavin adenine dinucleotide binding | $1.4 \times 10^{5}$ | $1 \times 10^{-6}$ | **0.00047*** |
| proteolysis | $7.6 \times 10^{5}$ | $2.1 \times 10^{-6}$ | **0.00096*** |
| lipid catabolic process | $1.7 \times 10^{5}$ | $2.2 \times 10^{-6}$ | **0.001*** |
| plasma membrane | $3.6 \times 10^{6}$ | $3.7 \times 10^{-6}$ | **0.002*** |
| cellular protein metabolic process | $2.5 \times 10^{5}$ | $5 \times 10^{-6}$ | **0.002*** |
| lyase activity | $2.2 \times 10^{5}$ | $5.6 \times 10^{-6}$ | **0.003*** |
| lysosomal lumen | $1.7 \times 10^{5}$ | $6.5 \times 10^{-6}$ | **0.003*** |
| extracellular matrix disassembly | $1.3 \times 10^{5}$ | $1.2 \times 10^{-5}$ | **0.005*** |
| organelle membrane | $1.4 \times 10^{5}$ | $1.3 \times 10^{-5}$ | **0.006*** |
| peptidase activity | $6.4 \times 10^{5}$ | $1.4 \times 10^{-5}$ | **0.006*** |
| endoplasmic reticulum lumen | $4.1 \times 10^{5}$ | $2 \times 10^{-5}$ | **0.009*** |
| mitochondrial matrix | $4.1 \times 10^{5}$ | $2.2 \times 10^{-5}$ | **0.010*** |
| iron ion binding | $1.9 \times 10^{5}$ | $2.3 \times 10^{-5}$ | **0.011*** |
| sodium ion transport | $2 \times 10^{5}$ | $2.9 \times 10^{-5}$ | **0.013*** |
| response to lipopolysaccharide | $2.3 \times 10^{5}$ | $3.4 \times 10^{-5}$ | **0.015*** |
| cytokine activity | $2.5 \times 10^{5}$ | $5.7 \times 10^{-5}$ | **0.026*** |
| calcium ion binding | $7.5 \times 10^{5}$ | $7.3 \times 10^{-5}$ | **0.033*** |
| viral entry into host cell | $1.3 \times 10^{5}$ | $7.7 \times 10^{-5}$ | **0.035*** |
| receptor-mediated endocytosis | $2 \times 10^{5}$ | 0.00011 | **0.049*** |
| collagen catabolic process | $1.3 \times 10^{5}$ | 0.00013 | 0.060 |
| carbohydrate binding | $2 \times 10^{5}$ | 0.00015 | 0.067 |
| cholesterol metabolic process | $1.5 \times 10^{5}$ | 0.00016 | 0.070 |
| platelet degranulation | $1.8 \times 10^{5}$ | 0.00017 | 0.075 |
| defense response to virus | $1.7 \times 10^{5}$ | 0.00017 | 0.075 |
| specific granule membrane | $1.5 \times 10^{5}$ | 0.00017 | 0.076 |
| basolateral plasma membrane | $2.5 \times 10^{5}$ | 0.00019 | 0.085 |
| cell adhesion | $6.4 \times 10^{5}$ | 0.00022 | 0.097 |
| acrosomal vesicle | $1.3 \times 10^{5}$ | 0.00023 | 0.101 |
| cell-matrix adhesion | $1.6 \times 10^{5}$ | 0.00024 | 0.104 |
| leukocyte migration | $2 \times 10^{5}$ | 0.00025 | 0.108 |
| cell surface receptor signaling pathway | $3.4 \times 10^{5}$ | 0.00025 | 0.109 |
| collagen binding | $1.3 \times 10^{5}$ | 0.00034 | 0.150 |
| heme binding | $1.4 \times 10^{5}$ | 0.00035 | 0.150 |
| endoplasmic reticulum | $1.5 \times 10^{6}$ | 0.00035 | 0.151 |
| chemotaxis | $1.7 \times 10^{5}$ | 0.00036 | 0.154 |
| transmembrane transporter activity | $1.6 \times 10^{5}$ | 0.00037 | 0.159 |
| carbohydrate metabolic process | $2.7 \times 10^{5}$ | 0.00037 | 0.159 |
| transmembrane transport | $6.6 \times 10^{5}$ | 0.00038 | 0.161 |
| extracellular matrix organization | $3 \times 10^{5}$ | 0.0006 | 0.256 |
| centriole | $1.6 \times 10^{5}$ | 0.001 | 0.440 |
| adaptive immune response | $1.7 \times 10^{5}$ | 0.001 | 0.462 |
| cilium | $3.4 \times 10^{5}$ | 0.001 | 0.493 |

Continued on next page

| Gene ontology | Mann-Whitney U | $p_v$ | $p_v^{adj}$ (* if < 0.05) |
|---|---|---|---|
| receptor activity | $2.9 \times 10^5$ | 0.001 | 0.542 |
| collagen trimer | $1.4 \times 10^5$ | 0.001 | 0.612 |
| extracellular matrix | $3.6 \times 10^5$ | 0.002 | 0.637 |
| protease binding | $1.4 \times 10^5$ | 0.002 | 0.637 |
| intracellular membrane-bounded organelle | $8.4 \times 10^5$ | 0.002 | 0.762 |
| lipid transport | $1.4 \times 10^5$ | 0.002 | 0.799 |

The sites detected under adaptation by mutation-selection codon models are enriched primarily with ontologies related to immune system processes (innate immune response, immune response, inflammatory response, defense response to virus, etc) and ontologies related to the external membrane (extracellular region, extracellular exosome, etc). The ontologies are similar to the sites detected by site-specific $\omega$-based model, although with stronger statistical support (lower $p_v^{adj}$). Moreover, sites detected under adaptation by mutation-selection codon models are also enriched with ontologies related to oxidoreductase activites, oxidation-reduction process and mitochondrion, which are not found by site-specific $\omega$-based model.

#### 2.4 Exclusive to site-specific mutation-selection model

For each gene, we computed the proportion of sites classified under an adaptive regime such that the lower bound for the posterior credibility interval of site-specific $\omega$ ($\alpha = 0.05$) is above the upper bound of the posterior credibility interval of site-specific $\omega_0$ ($\alpha = 0.05$), while the mean site-specific $\omega$ is below 1. Thus, these sites cannot be detected by site-specific mutation-selection model. Because this is a unilateral test ($\omega > \omega_0$) and the two posterior credibility interval are independent, the risk is $(\alpha/2)^2 = 0.025^2 = 6.25 \times 10^{-4}$ for each site. For each ontology, the proportion of sites under adaptation is compared between the set of genes sharing this given ontology and the rest of the genes with Mann-Whitney U test. $p_v^{adj}$ are corrected for multiple comparison (Holm–Bonferroni correction).

Table S4: Ontology enrichment with sites exclusive to mutation-selection model

| Gene ontology | Mann-Whitney U | $p_v$ | $p_v^{adj}$ (* if < 0.05) |
|---|---|---|---|
| oxidation-reduction process | $9.1 \times 10^5$ | $9.3 \times 10^{-26}$ | **$4.6 \times 10^{-23}$*** |
| extracellular exosome | $2.6 \times 10^6$ | $7.6 \times 10^{-25}$ | **$3.8 \times 10^{-22}$*** |
| oxidoreductase activity | $8.1 \times 10^5$ | $7.5 \times 10^{-24}$ | **$3.7 \times 10^{-21}$*** |
| lipid metabolic process | $7.7 \times 10^5$ | $1.6 \times 10^{-21}$ | **$8.1 \times 10^{-19}$*** |
| metabolic process | $6.2 \times 10^5$ | $7 \times 10^{-16}$ | **$3.4 \times 10^{-13}$*** |
| transmembrane transport | $7.5 \times 10^5$ | $1.1 \times 10^{-14}$ | **$5.3 \times 10^{-12}$*** |
| membrane | $4.8 \times 10^6$ | $2.2 \times 10^{-14}$ | **$1.1 \times 10^{-11}$*** |
| plasma membrane | $3.7 \times 10^6$ | $2.4 \times 10^{-14}$ | **$1.2 \times 10^{-11}$*** |
| integral component of membrane | $4.1 \times 10^6$ | $1 \times 10^{-13}$ | **$4.9 \times 10^{-11}$*** |
| integral component of plasma membrane | $1.5 \times 10^6$ | $6 \times 10^{-13}$ | **$2.9 \times 10^{-10}$*** |
| mitochondrion | $1.7 \times 10^6$ | $9.6 \times 10^{-13}$ | **$4.7 \times 10^{-10}$*** |
| catalytic activity | $6.9 \times 10^5$ | $1.1 \times 10^{-12}$ | **$5.4 \times 10^{-10}$*** |
| symporter activity | $2 \times 10^5$ | $5.3 \times 10^{-11}$ | **$2.5 \times 10^{-8}$*** |
| ion transport | $8 \times 10^5$ | $7.3 \times 10^{-11}$ | **$3.5 \times 10^{-8}$*** |
| transmembrane transporter activity | $2 \times 10^5$ | $7.3 \times 10^{-11}$ | **$3.5 \times 10^{-8}$*** |
| endoplasmic reticulum | $1.6 \times 10^6$ | $8 \times 10^{-11}$ | **$3.8 \times 10^{-8}$*** |
| fatty acid metabolic process | $2.6 \times 10^5$ | $1.2 \times 10^{-10}$ | **$5.7 \times 10^{-8}$*** |
| mitochondrial inner membrane | $4.9 \times 10^5$ | $5.6 \times 10^{-10}$ | **$2.7 \times 10^{-7}$*** |
| blood microparticle | $1.7 \times 10^5$ | $2.3 \times 10^{-9}$ | **$1.1 \times 10^{-6}$*** |
| lysosomal lumen | $1.8 \times 10^5$ | $5.9 \times 10^{-9}$ | **$2.8 \times 10^{-6}$*** |
| sodium ion transport | $2.2 \times 10^5$ | $1.2 \times 10^{-8}$ | **$5.7 \times 10^{-6}$*** |
| apical plasma membrane | $4.4 \times 10^5$ | $1 \times 10^{-7}$ | **$4.8 \times 10^{-5}$*** |
| flavin adenine dinucleotide binding | $1.4 \times 10^5$ | $2 \times 10^{-7}$ | **$9.4 \times 10^{-5}$*** |
| basolateral plasma membrane | $2.7 \times 10^5$ | $2.4 \times 10^{-7}$ | **0.00011*** |
| calcium ion binding | $7.7 \times 10^5$ | $2.4 \times 10^{-7}$ | **0.00011*** |
| receptor-mediated endocytosis | $2.1 \times 10^5$ | $3.5 \times 10^{-7}$ | **0.00016*** |
| neutrophil degranulation | $5.5 \times 10^5$ | $3.9 \times 10^{-7}$ | **0.00018*** |

Continued on next page

| Gene ontology | Mann-Whitney U | $p_{\mathrm{v}}$ | $p_{\mathrm{v}}^{\mathrm{adj}}$ (* if $< 0.05$) |
|---|---|---|---|
| peroxisome | $1.9 \times 10^5$ | $4.3 \times 10^{-7}$ | **0.0002*** |
| iron ion binding | $2 \times 10^5$ | $5.9 \times 10^{-7}$ | **0.00028*** |
| steroid metabolic process | $1.6 \times 10^5$ | $1.4 \times 10^{-6}$ | **0.00067*** |
| hydrolase activity | $1.7 \times 10^6$ | $2.8 \times 10^{-6}$ | **0.001*** |
| endoplasmic reticulum lumen | $4.1 \times 10^5$ | $2.8 \times 10^{-6}$ | **0.001*** |
| endoplasmic reticulum membrane | $1 \times 10^6$ | $4.2 \times 10^{-6}$ | **0.002*** |
| mitochondrial matrix | $4.2 \times 10^5$ | $6.9 \times 10^{-6}$ | **0.003*** |
| ligase activity | $2.2 \times 10^5$ | $8.1 \times 10^{-6}$ | **0.004*** |
| lyase activity | $2.1 \times 10^5$ | $8.3 \times 10^{-6}$ | **0.004*** |
| cellular protein metabolic process | $2.5 \times 10^5$ | $1.1 \times 10^{-5}$ | **0.005*** |
| intracellular membrane-bounded organelle | $8.7 \times 10^5$ | $1.4 \times 10^{-5}$ | **0.006*** |
| extracellular space | $1.4 \times 10^6$ | $1.5 \times 10^{-5}$ | **0.007*** |
| external side of plasma membrane | $2.6 \times 10^5$ | $2.2 \times 10^{-5}$ | **0.010*** |
| lipid transport | $1.5 \times 10^5$ | $3.2 \times 10^{-5}$ | **0.015*** |
| heme binding | $1.5 \times 10^5$ | $3.6 \times 10^{-5}$ | **0.016*** |
| organelle membrane | $1.3 \times 10^5$ | $5 \times 10^{-5}$ | **0.023*** |
| serine-type endopeptidase activity | $2.2 \times 10^5$ | $9.8 \times 10^{-5}$ | **0.045*** |
| endosome membrane | $2.9 \times 10^5$ | 0.00013 | 0.060 |
| extracellular matrix organization | $3.1 \times 10^5$ | 0.00013 | 0.060 |
| serine-type peptidase activity | $1.7 \times 10^5$ | 0.00017 | 0.075 |
| transporter activity | $1.9 \times 10^5$ | 0.00019 | 0.084 |
| lysosome | $4.8 \times 10^5$ | 0.00022 | 0.097 |
| phospholipid binding | $1.5 \times 10^5$ | 0.00031 | 0.136 |
| ATP binding | $1.6 \times 10^6$ | 0.00034 | 0.151 |
| peptidase activity | $6.2 \times 10^5$ | 0.00038 | 0.168 |
| carbohydrate metabolic process | $2.6 \times 10^5$ | 0.00041 | 0.184 |
| structural molecule activity | $2.1 \times 10^5$ | 0.00045 | 0.198 |
| myelin sheath | $1.7 \times 10^5$ | 0.00052 | 0.229 |
| platelet degranulation | $1.7 \times 10^5$ | 0.00058 | 0.256 |
| cholesterol metabolic process | $1.4 \times 10^5$ | 0.00062 | 0.272 |
| isomerase activity | $1.2 \times 10^5$ | 0.00067 | 0.292 |
| specific granule membrane | $1.4 \times 10^5$ | 0.00088 | 0.386 |
| chemotaxis | $1.7 \times 10^5$ | 0.00095 | 0.413 |
| proteolysis | $7.2 \times 10^5$ | 0.001 | 0.436 |
| sarcolemma | $1.4 \times 10^5$ | 0.001 | 0.470 |
| extracellular matrix | $3.6 \times 10^5$ | 0.001 | 0.543 |
| transferase activity | $1.8 \times 10^6$ | 0.001 | 0.563 |
| membrane raft | $2.8 \times 10^5$ | 0.002 | 0.676 |
| actin filament binding | $1.7 \times 10^5$ | 0.002 | 0.677 |
| cell surface | $6 \times 10^5$ | 0.002 | 0.697 |
| collagen catabolic process | $1.2 \times 10^5$ | 0.002 | 0.703 |
| calcium ion transmembrane transport | $1.5 \times 10^5$ | 0.002 | 0.727 |
| regulation of ion transmembrane transport | $1.8 \times 10^5$ | 0.002 | 0.915 |
| calcium ion transport | $1.6 \times 10^5$ | 0.002 | 0.981 |

The sites detected under adaptation solely by mutation-selection codon models (with $\omega < 1$) are enriched primarily with ontologies related to oxidoreductase activites, oxidation-reduction and mitochondrion process as well as ontologies related to the membrane (transmembrane transport, integral component of membrane, transmembrane transporter activity, etc).

### 3 Graphical abstract for the pipeline.

Figure S1:

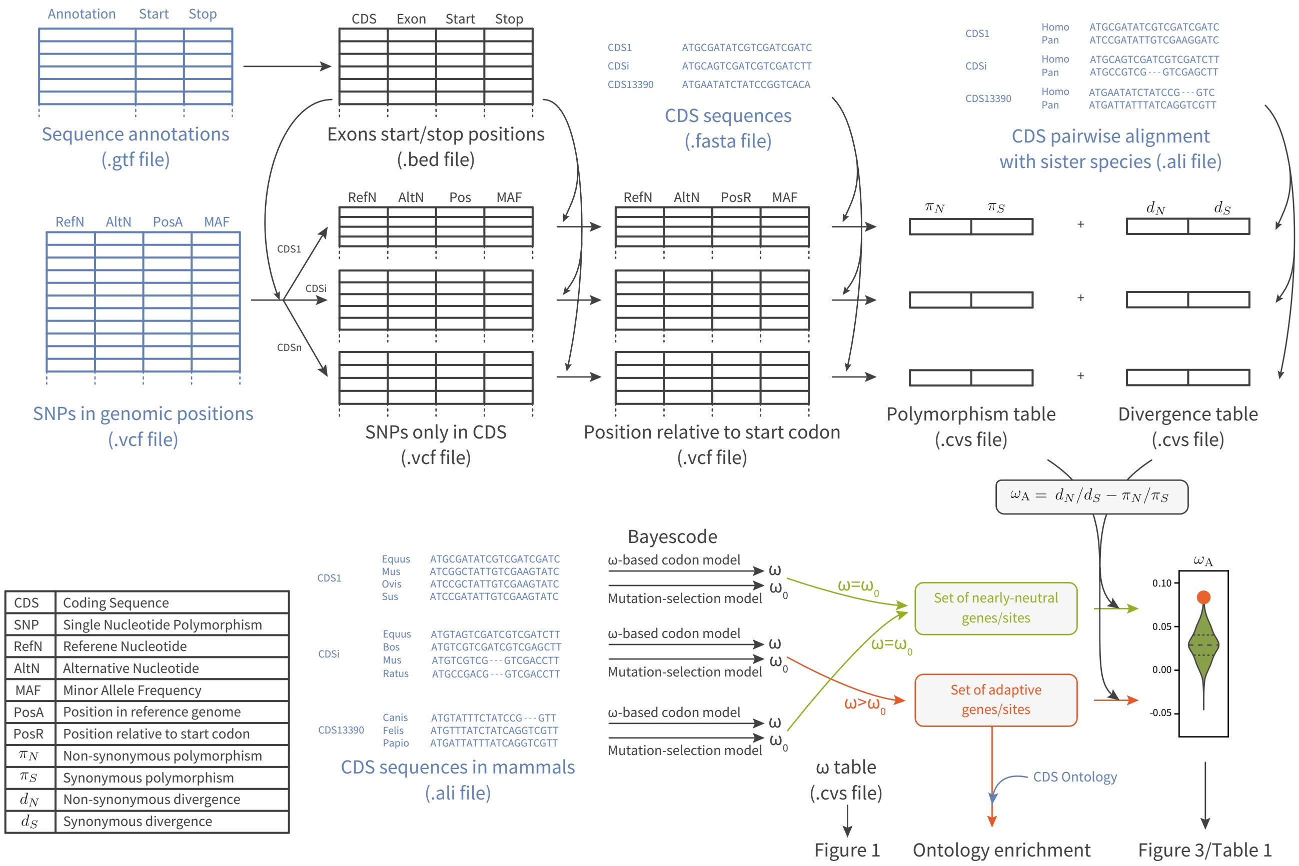


| | |
|---|---|
| CDS | Coding Sequence |
| SNP | Single Nucleotide Polymorphism |
| RefN | Referene Nucleotide |
| AltN | Alternative Nucleotide |
| MAF | Minor Allele Frequency |
| PosA | Position in reference genome |
| PosR | Position relative to start codon |
| $\pi_N$ | Non-synonymous polymorphism |
| $\pi_S$ | Synonymous polymorphism |
| $d_N$ | Non-synonymous divergence |
| $d_S$ | Synonymous divergence |

### 4 Rate of adaptation enrichment while controlling for $\omega$

$\omega$ is controlled to be the same in the nearly-neutral replicate and the adaptive set of genes, such as to alleviate the fact that genes classified as adaptive have a higher $\omega$ than genes classified as nearly-neutral, which could bias our comparison since $\omega_{\mathrm{A}}$ could simply be higher for genes with higher $\omega$.

Figure S2:

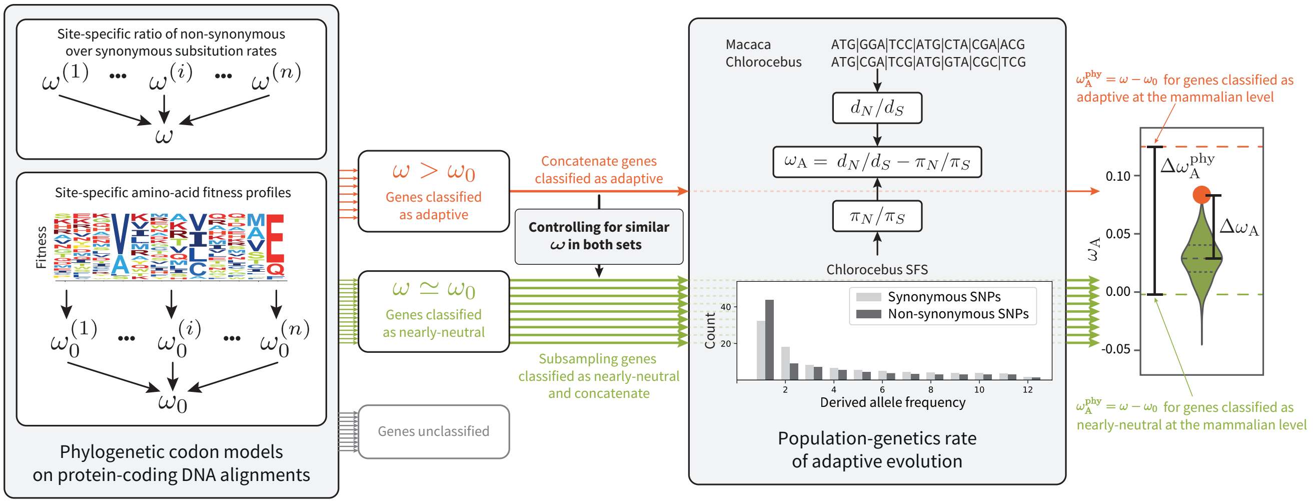


The random sampling is weighted to control for $\omega$ in the set of nearly-neutral genes/sites. First, a normal distribution is fitted to $\omega$ in both sets, and the probability density is called $f$ for the adaptive set and $g$ for the nearly-neutral set. Secondly, for each gene/site classified as nearly-neutral the weight is computed as the ratio $f(\omega)/g(\omega)$ for this specific gene/site. Sampling with this procedure produce a set of genes/sites classified as nearly-neutral with the same $\omega$ on average than the set of adaptive genes/sites.

##### 4.1 Mutation-selection codon model at gene level ($\alpha = 0.05$)

Figure S3:

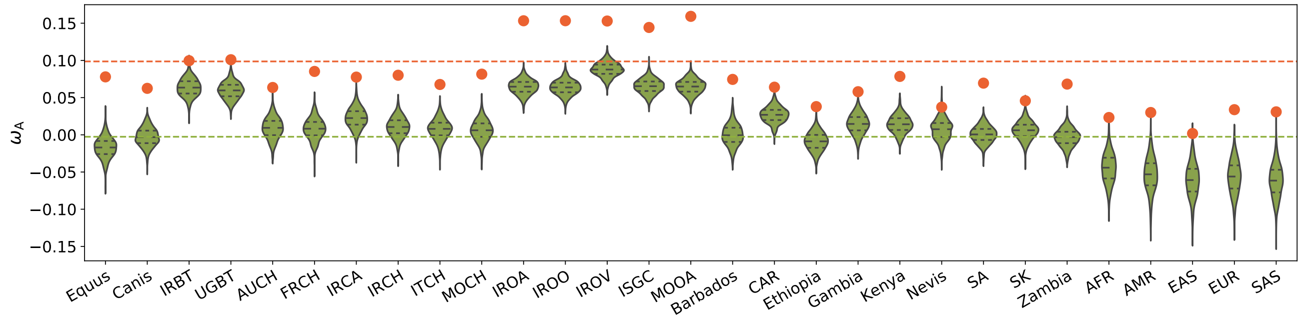


Table S5:

| Population | Species | $\omega_{\mathrm{A}}$ Adaptive | $\langle\omega_{\mathrm{A}}\rangle$ Nearly-neutral | $\Delta\omega_{\mathrm{A}}$ | $p_{\mathrm{v}}$ | $p_{\mathrm{v}}^{\mathrm{adj}}$ | $\frac{\Delta\omega_{\mathrm{A}}}{\omega_{\mathrm{A}}^{\mathrm{phy}}}$ | $\pi_{\mathrm{S}}$ |
|---|---|---|---|---|---|---|---|---|
| Diverse (Equus) | Equus caballus | 0.078 | −0.017 | 0.095 | 0.0 | **0.0*** | 0.952 | 0.002 |
| Diverse (Canis) | Canis familiaris | 0.062 | −0.003 | 0.065 | 0.0 | **0.0*** | 0.639 | 0.004 |
| Iran (IRBT) | Bos taurus | 0.100 | 0.064 | 0.036 | 0.002 | **0.006*** | 0.356 | 0.007 |
| Uganda (UGBT) | Bos taurus | 0.101 | 0.059 | 0.041 | 0.0 | **0.0*** | 0.412 | 0.008 |
| Australia (AUCH) | Capra hircus | 0.064 | 0.009 | 0.055 | 0.0 | **0.0*** | 0.541 | 0.003 |
| France (FRCH) | Capra hircus | 0.085 | 0.008 | 0.077 | 0.0 | **0.0*** | 0.764 | 0.002 |
| Iran (IRCA) | Capra aegagrus | 0.078 | 0.022 | 0.055 | 0.0 | **0.0*** | 0.544 | 0.003 |
| Iran (IRCH) | Capra hircus | 0.080 | 0.011 | 0.070 | 0.0 | **0.0*** | 0.688 | 0.004 |
| Italy (ITCH) | Capra hircus | 0.068 | 0.008 | 0.060 | 0.0 | **0.0*** | 0.594 | 0.003 |
| Morocco (MOCH) | Capra hircus | 0.082 | 0.006 | 0.076 | 0.0 | **0.0*** | 0.748 | 0.004 |
| Iran (IROA) | Ovis aries | 0.153 | 0.064 | 0.089 | 0.0 | **0.0*** | 0.878 | 0.007 |
| Iran (IROO) | Ovis orientalis | 0.153 | 0.064 | 0.090 | 0.0 | **0.0*** | 0.886 | 0.008 |
| Iran (IROV) | Ovis vignei | 0.153 | 0.088 | 0.065 | 0.0 | **0.0*** | 0.642 | 0.005 |
| Various (ISGC) | Ovis aries | 0.144 | 0.065 | 0.079 | 0.0 | **0.0*** | 0.780 | 0.008 |
| Morocco (MOOA) | Ovis aries | 0.159 | 0.065 | 0.095 | 0.0 | **0.0*** | 0.934 | 0.007 |
| Barbados | Chlorocebus sabaeus | 0.074 | −0.0002 | 0.075 | 0.0 | **0.0*** | 0.737 | 0.003 |
| Central African Republic (CAR) | Chlorocebus sabaeus | 0.064 | 0.027 | 0.037 | 0.0 | **0.0*** | 0.368 | 0.006 |
| Ethiopia | Chlorocebus sabaeus | 0.038 | −0.009 | 0.047 | 0.0 | **0.0*** | 0.461 | 0.005 |
| Gambia | Chlorocebus sabaeus | 0.058 | 0.014 | 0.043 | 0.0 | **0.0*** | 0.429 | 0.005 |
| Kenya | Chlorocebus sabaeus | 0.079 | 0.014 | 0.064 | 0.0 | **0.0*** | 0.634 | 0.004 |
| Nevis | Chlorocebus sabaeus | 0.037 | 0.007 | 0.030 | 0.020 | **0.020*** | 0.300 | 0.003 |
| South Africa (SA) | Chlorocebus sabaeus | 0.069 | 0.00053 | 0.069 | 0.0 | **0.0*** | 0.680 | 0.006 |
| Saint Kitts (SK) | Chlorocebus sabaeus | 0.046 | 0.006 | 0.040 | 0.001 | **0.004*** | 0.393 | 0.004 |
| Zambia | Chlorocebus sabaeus | 0.068 | −0.004 | 0.072 | 0.0 | **0.0*** | 0.709 | 0.006 |
| African (AFR) | Homo sapiens | 0.023 | −0.045 | 0.068 | 0.0 | **0.0*** | 0.672 | 0.002 |
| Ad Mixed American (AMR) | Homo sapiens | 0.030 | −0.054 | 0.083 | 0.0 | **0.0*** | 0.825 | 0.002 |
| East Asian (EAS) | Homo sapiens | 0.002 | −0.061 | 0.063 | 0.002 | **0.006*** | 0.620 | 0.002 |
| European (EUR) | Homo sapiens | 0.034 | −0.057 | 0.091 | 0.0 | **0.0*** | 0.896 | 0.002 |
| South Asian (SAS) | Homo sapiens | 0.031 | −0.062 | 0.093 | 0.0 | **0.0*** | 0.917 | 0.002 |

Figure S4:

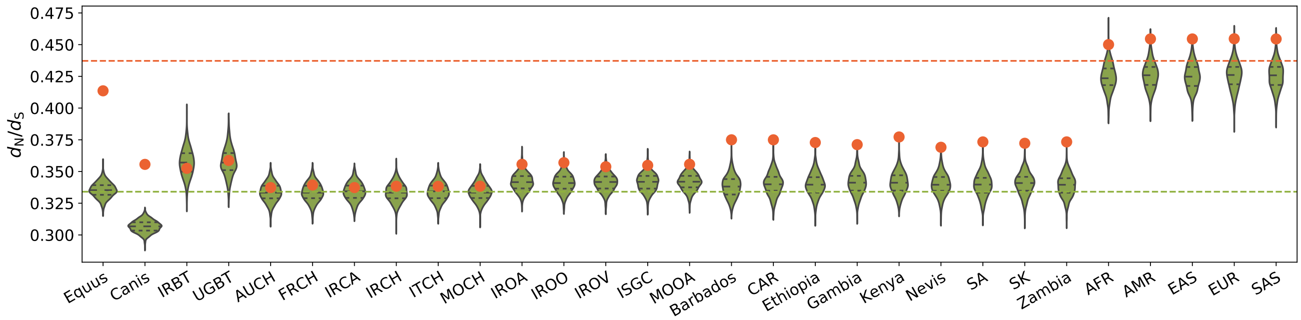


Table S6:

| Population | Species | $d_{\mathrm{N}}/d_{\mathrm{S}}$ Adaptive | $\langle d_{\mathrm{N}}/d_{\mathrm{S}} \rangle$ Nearly-neutral | $p_{\mathrm{v}}$ | $p_{\mathrm{v}}^{\mathrm{adj}}$ | $\pi_{\mathrm{S}}$ |
|---|---|---|---|---|---|---|
| Diverse (Equus) | Equus caballus | 0.414 | 0.335 | 0.0 | **0.0*** | 0.002 |
| Diverse (Canis) | Canis familiaris | 0.356 | 0.307 | 0.0 | **0.0*** | 0.004 |
| Iran (IRBT) | Bos taurus | 0.352 | 0.357 | 0.657 | 1.000 | 0.007 |
| Uganda (UGBT) | Bos taurus | 0.359 | 0.358 | 0.432 | 1.000 | 0.008 |
| Australia (AUCH) | Capra hircus | 0.337 | 0.334 | 0.312 | 1.000 | 0.003 |
| France (FRCH) | Capra hircus | 0.339 | 0.334 | 0.223 | 1.000 | 0.002 |
| Iran (IRCA) | Capra aegagrus | 0.337 | 0.334 | 0.338 | 1.000 | 0.003 |
| Iran (IRCH) | Capra hircus | 0.338 | 0.334 | 0.257 | 1.000 | 0.004 |
| Italy (ITCH) | Capra hircus | 0.338 | 0.334 | 0.278 | 1.000 | 0.003 |
| Morocco (MOCH) | Capra hircus | 0.338 | 0.334 | 0.252 | 1.000 | 0.004 |
| Iran (IROA) | Ovis aries | 0.356 | 0.342 | 0.021 | 0.231 | 0.007 |
| Iran (IROO) | Ovis orientalis | 0.357 | 0.341 | 0.013 | 0.169 | 0.008 |
| Iran (IROV) | Ovis vignei | 0.354 | 0.342 | 0.037 | 0.333 | 0.005 |
| Various (ISGC) | Ovis aries | 0.355 | 0.342 | 0.025 | 0.250 | 0.008 |
| Morocco (MOOA) | Ovis aries | 0.356 | 0.342 | 0.017 | 0.204 | 0.007 |
| Barbados | Chlorocebus sabaeus | 0.375 | 0.338 | 0.0 | **0.0*** | 0.003 |
| Central African Republic (CAR) | Chlorocebus sabaeus | 0.375 | 0.340 | 0.0 | **0.0*** | 0.006 |
| Ethiopia | Chlorocebus sabaeus | 0.373 | 0.340 | 0.0 | **0.0*** | 0.005 |
| Gambia | Chlorocebus sabaeus | 0.371 | 0.341 | 0.0 | **0.0*** | 0.005 |
| Kenya | Chlorocebus sabaeus | 0.377 | 0.341 | 0.0 | **0.0*** | 0.004 |
| Nevis | Chlorocebus sabaeus | 0.369 | 0.340 | 0.0 | **0.0*** | 0.003 |
| South Africa (SA) | Chlorocebus sabaeus | 0.373 | 0.339 | 0.0 | **0.0*** | 0.006 |
| Saint Kitts (SK) | Chlorocebus sabaeus | 0.372 | 0.340 | 0.0 | **0.0*** | 0.004 |
| Zambia | Chlorocebus sabaeus | 0.373 | 0.339 | 0.0 | **0.0*** | 0.006 |
| African (AFR) | Homo sapiens | 0.450 | 0.424 | 0.005 | 0.075 | 0.002 |
| Ad Mixed American (AMR) | Homo sapiens | 0.455 | 0.426 | 0.002 | **0.034*** | 0.002 |
| East Asian (EAS) | Homo sapiens | 0.454 | 0.425 | 0.0 | **0.0*** | 0.002 |
| European (EUR) | Homo sapiens | 0.455 | 0.426 | 0.003 | **0.048*** | 0.002 |
| South Asian (SAS) | Homo sapiens | 0.454 | 0.425 | 0.005 | 0.075 | 0.002 |

In *Bos*, *Capras* and *Ovis*, $d_N/d_S$ computed for the set of genes under adaptation at the phylogenetic scale (red, $1 > \omega > \omega_0$) is not higher than for the set of genes under a nearly-neutral regime (green, $1 > \omega \simeq \omega_0$), showing that the sampling procedure controlling for $\omega$ at the phylogenetic scale is valid. However, even though $d_N/d_S$ is not higher (genes are not faster overall), the rate of adaptation at the population-genetic scale ($\omega_{\mathrm{A}}$) for the set of genes supposedly under adaptation is higher than for the set of nearly-neutral genes in all populations. Altogether, these genes have a higher rate of adaptation, while not evolving faster than there nearly-neutral counterpart, showing that the rate of adaptation computed at the phylogenetic scale is able to detect genes under pervasive adaptation.

##### 4.2 Mutation-selection codon model at site level ($\alpha = 0.05$)

Figure S5:

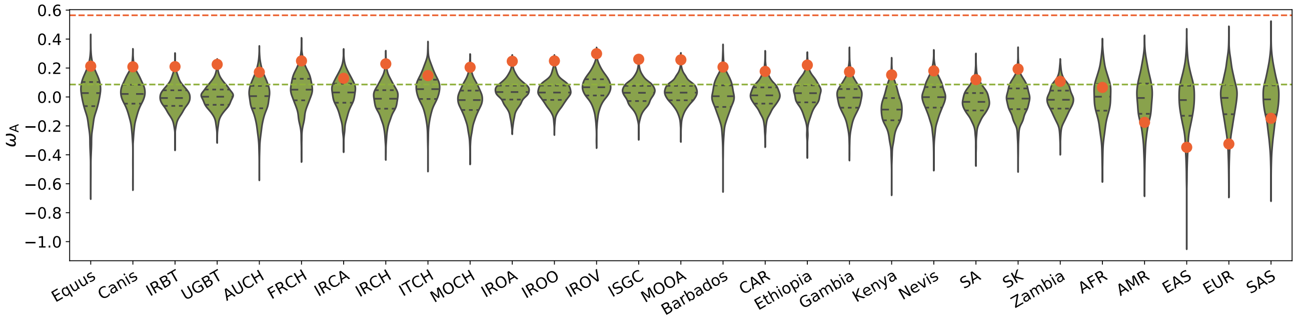


Table S7:

| Population | Species | $\omega_A$ Adaptive | $\langle\omega_A\rangle$ Nearly-neutral | $\Delta\omega_A$ | $p_v$ | $p_v^{adj}$ | $\frac{\Delta\omega_A}{\omega_A^{phy}}$ | $\pi_S$ |
|---|---|---|---|---|---|---|---|---|
| Diverse (Equus) | Equus caballus | 0.213 | 0.010 | 0.203 | 0.034 | 0.374 | 0.425 | 0.002 |
| Diverse (Canis) | Canis familiaris | 0.208 | 0.013 | 0.195 | 0.013 | 0.234 | 0.407 | 0.004 |
| Iran (IRBT) | Bos taurus | 0.210 | −0.007 | 0.217 | 0.005 | 0.110 | 0.454 | 0.007 |
| Uganda (UGBT) | Bos taurus | 0.225 | −0.002 | 0.227 | 0.0 | **0.0*** | 0.476 | 0.008 |
| Australia (AUCH) | Capra hircus | 0.171 | −0.006 | 0.177 | 0.056 | 0.504 | 0.369 | 0.003 |
| France (FRCH) | Capra hircus | 0.249 | 0.046 | 0.204 | 0.025 | 0.336 | 0.425 | 0.002 |
| Iran (IRCA) | Capra aegagrus | 0.129 | 0.023 | 0.106 | 0.149 | 1.000 | 0.221 | 0.003 |
| Iran (IRCH) | Capra hircus | 0.229 | −0.020 | 0.249 | 0.004 | 0.092 | 0.519 | 0.004 |
| Italy (ITCH) | Capra hircus | 0.148 | 0.048 | 0.099 | 0.170 | 1.000 | 0.207 | 0.003 |
| Morocco (MOCH) | Capra hircus | 0.205 | −0.026 | 0.231 | 0.009 | 0.171 | 0.482 | 0.004 |
| Iran (IROA) | Ovis aries | 0.247 | 0.028 | 0.219 | 0.002 | **0.048*** | 0.456 | 0.007 |
| Iran (IROO) | Ovis orientalis | 0.249 | 0.029 | 0.220 | 0.001 | **0.026*** | 0.458 | 0.008 |
| Iran (IROV) | Ovis vignei | 0.299 | 0.063 | 0.236 | 0.0 | **0.0*** | 0.493 | 0.005 |
| Various (ISGC) | Ovis aries | 0.261 | 0.024 | 0.237 | 0.0 | **0.0*** | 0.493 | 0.008 |
| Morocco (MOOA) | Ovis aries | 0.257 | 0.026 | 0.231 | 0.001 | **0.026*** | 0.481 | 0.007 |
| Barbados | Chlorocebus sabaeus | 0.207 | 0.00044 | 0.206 | 0.017 | 0.272 | 0.432 | 0.003 |
| Central African Republic (CAR) | Chlorocebus sabaeus | 0.177 | 0.008 | 0.168 | 0.017 | 0.272 | 0.353 | 0.006 |
| Ethiopia | Chlorocebus sabaeus | 0.220 | 0.018 | 0.202 | 0.008 | 0.160 | 0.423 | 0.005 |
| Gambia | Chlorocebus sabaeus | 0.174 | −0.011 | 0.185 | 0.024 | 0.336 | 0.387 | 0.005 |
| Kenya | Chlorocebus sabaeus | 0.152 | −0.088 | 0.240 | 0.006 | 0.126 | 0.503 | 0.004 |
| Nevis | Chlorocebus sabaeus | 0.180 | −0.008 | 0.189 | 0.031 | 0.372 | 0.395 | 0.003 |
| South Africa (SA) | Chlorocebus sabaeus | 0.120 | −0.036 | 0.155 | 0.034 | 0.374 | 0.325 | 0.006 |
| Saint Kitts (SK) | Chlorocebus sabaeus | 0.193 | −0.018 | 0.211 | 0.013 | 0.234 | 0.441 | 0.004 |
| Zambia | Chlorocebus sabaeus | 0.108 | −0.021 | 0.129 | 0.067 | 0.536 | 0.269 | 0.006 |
| African (AFR) | Homo sapiens | 0.066 | −0.009 | 0.075 | 0.325 | 1.000 | 0.157 | 0.002 |
| Ad Mixed American (AMR) | Homo sapiens | −0.175 | −0.019 | −0.157 | 0.849 | 1.000 | −0.328 | 0.002 |
| East Asian (EAS) | Homo sapiens | −0.348 | −0.036 | −0.312 | 0.953 | 1.000 | −0.653 | 0.002 |
| European (EUR) | Homo sapiens | −0.325 | −0.021 | −0.304 | 0.965 | 1.000 | −0.636 | 0.002 |
| South Asian (SAS) | Homo sapiens | −0.148 | −0.025 | −0.122 | 0.798 | 1.000 | −0.256 | 0.002 |

Figure S6:

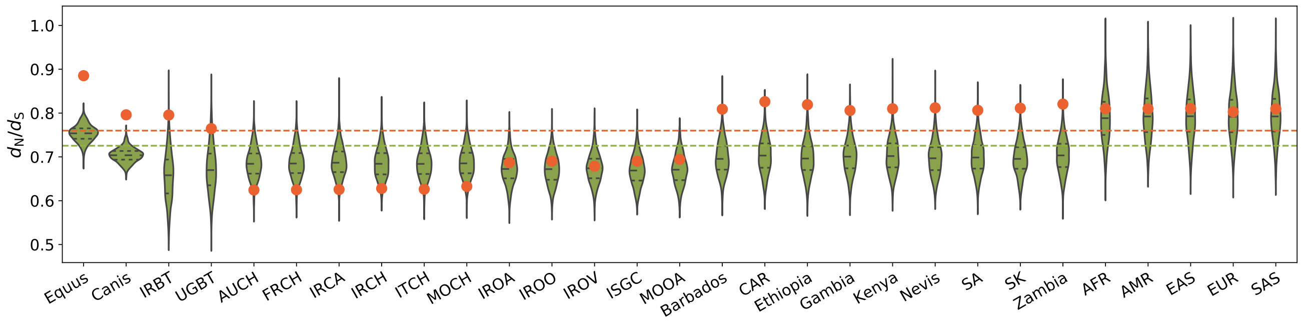


Table S8:

| Population | Species | $d_N/d_S$ Adaptive | $\langle d_N/d_S \rangle$ Nearly-neutral | $p_v$ | $p_v^{adj}$ | $\pi_S$ |
|---|---|---|---|---|---|---|
| Diverse (Equus) | Equus caballus | 0.885 | 0.754 | 0.0 | **0.0*** | 0.002 |
| Diverse (Canis) | Canis familiaris | 0.796 | 0.704 | 0.0 | **0.0*** | 0.004 |
| Iran (IRBT) | Bos taurus | 0.796 | 0.658 | 0.008 | 0.152 | 0.007 |
| Uganda (UGBT) | Bos taurus | 0.765 | 0.672 | 0.046 | 0.782 | 0.008 |
| Australia (AUCH) | Capra hircus | 0.624 | 0.686 | 0.972 | 1.000 | 0.003 |
| France (FRCH) | Capra hircus | 0.625 | 0.687 | 0.977 | 1.000 | 0.002 |
| Iran (IRCA) | Capra aegagrus | 0.626 | 0.690 | 0.977 | 1.000 | 0.003 |
| Iran (IRCH) | Capra hircus | 0.628 | 0.685 | 0.968 | 1.000 | 0.004 |
| Italy (ITCH) | Capra hircus | 0.626 | 0.686 | 0.970 | 1.000 | 0.003 |
| Morocco (MOCH) | Capra hircus | 0.633 | 0.686 | 0.948 | 1.000 | 0.004 |
| Iran (IROA) | Ovis aries | 0.687 | 0.672 | 0.331 | 1.000 | 0.007 |
| Iran (IROO) | Ovis orientalis | 0.690 | 0.672 | 0.298 | 1.000 | 0.008 |
| Iran (IROV) | Ovis vignei | 0.679 | 0.674 | 0.431 | 1.000 | 0.005 |
| Various (ISGC) | Ovis aries | 0.690 | 0.670 | 0.273 | 1.000 | 0.008 |
| Morocco (MOOA) | Ovis aries | 0.694 | 0.671 | 0.245 | 1.000 | 0.007 |
| Barbados | Chlorocebus sabaeus | 0.809 | 0.698 | 0.005 | 0.115 | 0.003 |
| Central African Republic (CAR) | Chlorocebus sabaeus | 0.826 | 0.704 | 0.003 | 0.081 | 0.006 |
| Ethiopia | Chlorocebus sabaeus | 0.819 | 0.697 | 0.003 | 0.081 | 0.005 |
| Gambia | Chlorocebus sabaeus | 0.806 | 0.702 | 0.012 | 0.216 | 0.005 |
| Kenya | Chlorocebus sabaeus | 0.810 | 0.703 | 0.006 | 0.126 | 0.004 |
| Nevis | Chlorocebus sabaeus | 0.812 | 0.697 | 0.004 | 0.100 | 0.003 |
| South Africa (SA) | Chlorocebus sabaeus | 0.806 | 0.701 | 0.007 | 0.140 | 0.006 |
| Saint Kitts (SK) | Chlorocebus sabaeus | 0.811 | 0.697 | 0.004 | 0.100 | 0.004 |
| Zambia | Chlorocebus sabaeus | 0.821 | 0.704 | 0.005 | 0.115 | 0.006 |
| African (AFR) | Homo sapiens | 0.810 | 0.790 | 0.351 | 1.000 | 0.002 |
| Ad Mixed American (AMR) | Homo sapiens | 0.810 | 0.796 | 0.378 | 1.000 | 0.002 |
| East Asian (EAS) | Homo sapiens | 0.810 | 0.794 | 0.382 | 1.000 | 0.002 |
| European (EUR) | Homo sapiens | 0.803 | 0.794 | 0.432 | 1.000 | 0.002 |
| South Asian (SAS) | Homo sapiens | 0.810 | 0.795 | 0.383 | 1.000 | 0.002 |

In *Bos*, *Capras*, *Ovis*, *Chlorocebus* and *Homo*, $d_N/d_S$ computed for the set of sites under adaptation (red, $1 > \omega > \omega_0$) at the phylogenetic scale is not higher than for the set of sites under a nearly-neutral regime (green, $1 > \omega \simeq \omega_0$), showing that the sampling procedure controlling for $\omega$ at the phylogenetic scale is valid. However, even though $d_N/d_S$ is not higher (sites are not faster overall), the rate of adaptation at the population-genetic scale ($\omega_A$) for the set of sites supposedly under adaptation is higher than for the set of nearly-neutral sites in *Bos* and *Ovis*. Altogether, these sites have a higher rate of adaptation, while not evolving faster than there nearly-neutral counterpart, showing that the rate of adaptation computed at the phylogenetic scale is able to detect sites under pervasive adaptation, even though these sites are with $\omega < 1$ and thus cannot be detected by site-specific $\omega$-based codon models.

### 5 Rate of adaptation enrichment with $\alpha = 0.005$

For each protein-coding DNA alignment, the Monte-Carlo Markov-Chain (MCMC) is run during 2000 points using the BayesCode software, after a burn-in of 1000 points. The mean of $\omega$ and $\omega_0$ are computed across the MCMC (after burn-in), as well as the 99.5% posterior credibility interval ($\alpha = 0.005$) for each gene and site, which is more stringent than the 95% interval ($\alpha = 0.05$) as shown in the main manuscript. Genes and sites classified under an adaptive regime (in red) are rejecting the nearly-neutral assumption such that a lower bound for the posterior credibility interval of $\omega$ is above the upper bound of the posterior credibility interval of $\omega_0$, meaning $\omega > \omega_0$. Genes and sites are classified under a nearly-neutral regime (in green) if the average $\omega$ is within the posterior credibility interval of the $\omega_0$, and respectively the average $\omega_0$ is also within the posterior credibility interval of $\omega$, meaning $\omega = \omega_0$. Genes and sites that do not fall in any of these categories are considered unclassified.

#### 5.1 Scatterplot with $\alpha = 0.005$

Figure S7:

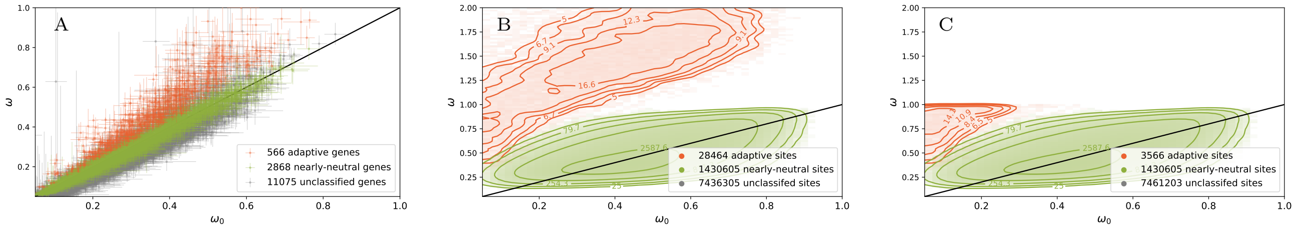


$\omega$ estimated by the site model against $\omega_0$ calculated by the mutation-selection model. Scatter plot of $14,509$ genes in panel A, with 99.5% posterior credibility interval ($\alpha = 0.005$). Density plot of sites in panel B and C. Genes or sites are then classified whether they detected as adaptive ($\omega > \omega_0$ in red) or nearly-neutral ($\omega \simeq \omega_0$ in green). In panel C, the set of sites detected exclusively by mutation-selection codon models have a mean $\omega < 1$.

#### 5.2 Mutation-selection codon model at gene level ($\alpha = 0.005$)

Figure S8:

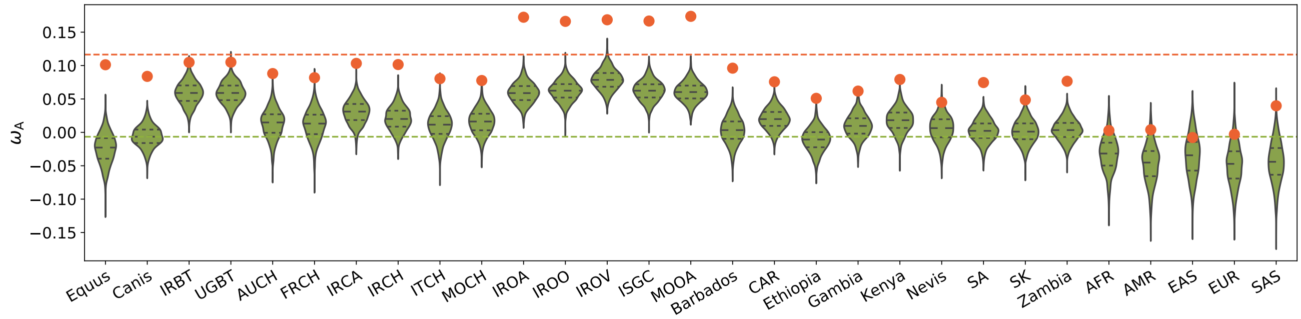


Table S9:

| Population | Species | $\omega_A$ Adaptive | $\langle\omega_A\rangle$ Nearly-neutral | $\Delta\omega_A$ | $p_v$ | $p_v^{adj}$ | $\frac{\Delta\omega_A}{\omega_A^{phy}}$ | $\pi_S$ |
|---|---|---|---|---|---|---|---|---|
| Diverse (Equus) | Equus caballus | 0.101 | −0.024 | 0.125 | 0.0 | **0.0*** | 1.029 | 0.002 |
| Diverse (Canis) | Canis familiaris | 0.084 | −0.006 | 0.090 | 0.0 | **0.0*** | 0.722 | 0.004 |
| Iran (IRBT) | Bos taurus | 0.105 | 0.058 | 0.046 | 0.002 | **0.014*** | 0.381 | 0.008 |
| Uganda (UGBT) | Bos taurus | 0.105 | 0.059 | 0.046 | 0.001 | **0.010*** | 0.382 | 0.008 |
| Australia (AUCH) | Capra hircus | 0.088 | 0.013 | 0.075 | 0.0 | **0.0*** | 0.611 | 0.003 |
| France (FRCH) | Capra hircus | 0.082 | 0.012 | 0.070 | 0.001 | **0.010*** | 0.573 | 0.003 |
| Iran (IRCA) | Capra aegagrus | 0.103 | 0.031 | 0.073 | 0.0 | **0.0*** | 0.593 | 0.004 |
| Iran (IRCH) | Capra hircus | 0.102 | 0.020 | 0.081 | 0.0 | **0.0*** | 0.662 | 0.004 |
| Italy (ITCH) | Capra hircus | 0.080 | 0.010 | 0.070 | 0.0 | **0.0*** | 0.571 | 0.003 |
| Morocco (MOCH) | Capra hircus | 0.078 | 0.016 | 0.062 | 0.0 | **0.0*** | 0.504 | 0.004 |
| Iran (IROA) | Ovis aries | 0.172 | 0.059 | 0.113 | 0.0 | **0.0*** | 0.919 | 0.007 |
| Iran (IROO) | Ovis orientalis | 0.166 | 0.062 | 0.104 | 0.0 | **0.0*** | 0.841 | 0.009 |
| Iran (IROV) | Ovis vignei | 0.169 | 0.079 | 0.090 | 0.0 | **0.0*** | 0.726 | 0.005 |
| Various (ISGC) | Ovis aries | 0.167 | 0.062 | 0.105 | 0.0 | **0.0*** | 0.847 | 0.008 |
| Morocco (MOOA) | Ovis aries | 0.174 | 0.060 | 0.113 | 0.0 | **0.0*** | 0.916 | 0.008 |
| Barbados | Chlorocebus sabaeus | 0.096 | 0.003 | 0.093 | 0.0 | **0.0*** | 0.755 | 0.003 |
| Central African Republic (CAR) | Chlorocebus sabaeus | 0.076 | 0.020 | 0.056 | 0.0 | **0.0*** | 0.451 | 0.006 |
| Ethiopia | Chlorocebus sabaeus | 0.051 | −0.011 | 0.062 | 0.0 | **0.0*** | 0.504 | 0.005 |
| Gambia | Chlorocebus sabaeus | 0.062 | 0.010 | 0.052 | 0.0 | **0.0*** | 0.423 | 0.005 |
| Kenya | Chlorocebus sabaeus | 0.079 | 0.018 | 0.061 | 0.0 | **0.0*** | 0.497 | 0.004 |
| Nevis | Chlorocebus sabaeus | 0.045 | 0.006 | 0.039 | 0.014 | 0.070 | 0.319 | 0.003 |
| South Africa (SA) | Chlorocebus sabaeus | 0.075 | 0.002 | 0.072 | 0.0 | **0.0*** | 0.587 | 0.006 |
| Saint Kitts (SK) | Chlorocebus sabaeus | 0.049 | 0.001 | 0.047 | 0.002 | **0.014*** | 0.384 | 0.004 |
| Zambia | Chlorocebus sabaeus | 0.077 | 0.004 | 0.073 | 0.0 | **0.0*** | 0.589 | 0.006 |
| African (AFR) | Homo sapiens | 0.003 | −0.033 | 0.036 | 0.071 | 0.174 | 0.289 | 0.002 |
| Ad Mixed American (AMR) | Homo sapiens | 0.004 | −0.047 | 0.051 | 0.033 | 0.132 | 0.415 | 0.002 |
| East Asian (EAS) | Homo sapiens | −0.008 | −0.036 | 0.028 | 0.186 | 0.186 | 0.227 | 0.002 |
| European (EUR) | Homo sapiens | −0.003 | −0.049 | 0.046 | 0.058 | 0.174 | 0.376 | 0.002 |
| South Asian (SAS) | Homo sapiens | 0.040 | −0.045 | 0.084 | 0.001 | **0.010*** | 0.688 | 0.002 |

#### 5.3 Mutation-selection codon model at site level ($\alpha = 0.005$)

Figure S9:

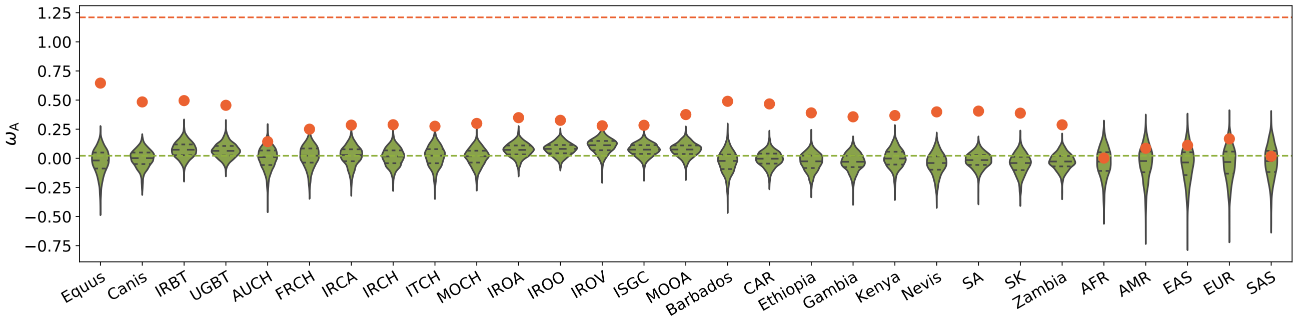


Table S10:

| Population | Species | $\omega_A$ Adaptive | $\langle\omega_A\rangle$ Nearly-neutral | $\Delta\omega_A$ | $p_v$ | $p_v^{adj}$ | $\frac{\Delta\omega_A}{\omega_A^{phy}}$ | $\pi_S$ |
|---|---|---|---|---|---|---|---|---|
| Diverse (Equus) | Equus caballus | 0.646 | −0.028 | 0.674 | 0.0 | **0.0*** | 0.567 | 0.002 |
| Diverse (Canis) | Canis familiaris | 0.484 | −0.003 | 0.487 | 0.0 | **0.0*** | 0.406 | 0.004 |
| Iran (IRBT) | Bos taurus | 0.495 | 0.072 | 0.423 | 0.0 | **0.0*** | 0.356 | 0.008 |
| Uganda (UGBT) | Bos taurus | 0.455 | 0.065 | 0.390 | 0.0 | **0.0*** | 0.328 | 0.008 |
| Australia (AUCH) | Capra hircus | 0.143 | 0.001 | 0.142 | 0.043 | 0.258 | 0.119 | 0.003 |
| France (FRCH) | Capra hircus | 0.250 | 0.017 | 0.233 | 0.0 | **0.0*** | 0.196 | 0.003 |
| Iran (IRCA) | Capra aegagrus | 0.285 | 0.021 | 0.264 | 0.0 | **0.0*** | 0.222 | 0.004 |
| Iran (IRCH) | Capra hircus | 0.288 | 0.012 | 0.276 | 0.0 | **0.0*** | 0.233 | 0.004 |
| Italy (ITCH) | Capra hircus | 0.275 | 0.015 | 0.260 | 0.0 | **0.0*** | 0.219 | 0.003 |
| Morocco (MOCH) | Capra hircus | 0.299 | 0.012 | 0.287 | 0.0 | **0.0*** | 0.242 | 0.004 |
| Iran (IROA) | Ovis aries | 0.349 | 0.070 | 0.279 | 0.0 | **0.0*** | 0.235 | 0.007 |
| Iran (IROO) | Ovis orientalis | 0.326 | 0.078 | 0.248 | 0.0 | **0.0*** | 0.209 | 0.009 |
| Iran (IROV) | Ovis vignei | 0.279 | 0.106 | 0.174 | 0.0 | **0.0*** | 0.146 | 0.005 |
| Various (ISGC) | Ovis aries | 0.284 | 0.072 | 0.211 | 0.0 | **0.0*** | 0.178 | 0.008 |
| Morocco (MOOA) | Ovis aries | 0.376 | 0.074 | 0.302 | 0.0 | **0.0*** | 0.254 | 0.008 |
| Barbados | Chlorocebus sabaeus | 0.490 | −0.031 | 0.521 | 0.0 | **0.0*** | 0.438 | 0.003 |
| Central African Republic (CAR) | Chlorocebus sabaeus | 0.467 | −0.006 | 0.473 | 0.0 | **0.0*** | 0.398 | 0.006 |
| Ethiopia | Chlorocebus sabaeus | 0.390 | −0.032 | 0.422 | 0.0 | **0.0*** | 0.355 | 0.005 |
| Gambia | Chlorocebus sabaeus | 0.356 | −0.031 | 0.388 | 0.0 | **0.0*** | 0.326 | 0.005 |
| Kenya | Chlorocebus sabaeus | 0.367 | −0.00077 | 0.368 | 0.0 | **0.0*** | 0.310 | 0.004 |
| Nevis | Chlorocebus sabaeus | 0.399 | −0.045 | 0.444 | 0.0 | **0.0*** | 0.373 | 0.003 |
| South Africa (SA) | Chlorocebus sabaeus | 0.405 | −0.017 | 0.422 | 0.0 | **0.0*** | 0.355 | 0.006 |
| Saint Kitts (SK) | Chlorocebus sabaeus | 0.388 | −0.048 | 0.436 | 0.0 | **0.0*** | 0.366 | 0.004 |
| Zambia | Chlorocebus sabaeus | 0.288 | −0.028 | 0.315 | 0.0 | **0.0*** | 0.265 | 0.006 |
| African (AFR) | Homo sapiens | 0.002 | −0.034 | 0.036 | 0.423 | 0.794 | 0.030 | 0.002 |
| Ad Mixed American (AMR) | Homo sapiens | 0.086 | −0.041 | 0.127 | 0.170 | 0.510 | 0.107 | 0.002 |
| East Asian (EAS) | Homo sapiens | 0.111 | −0.052 | 0.164 | 0.107 | 0.428 | 0.138 | 0.002 |
| European (EUR) | Homo sapiens | 0.167 | −0.045 | 0.212 | 0.059 | 0.295 | 0.178 | 0.002 |
| South Asian (SAS) | Homo sapiens | 0.017 | −0.031 | 0.048 | 0.397 | 0.794 | 0.040 | 0.002 |

With a threshold of $\alpha = 0.005$ more stringent than as shown in the main manuscript ($\alpha = 0.05$), the rate of false positive is mechanically lower both at the site and gene level. However, the statistical power to test for enrichment of adaptation is not necessarily higher since we are working with fewer data. We thus obtain high values of $\omega_A$ for the set of genes and sites supposedly under adaptation at the phylogenetic scale, which also leads to higher $\Delta\omega_A$ when compared to the the set of genes and sites supposedly under a nearly-neutral regime. Since we have more variance due to less data (violin plots are more extended for the set of nearly-neutral replicates), this higher value of the statistic $\Delta\omega_A$ does not translate to lower $p_v^{adj}$. We thus have to find a compromise between having large enough dataset to perform reliable computation of $\omega_A$ and having a very low rate of false positive for sites and genes under adaptation at the phylogenetic scale. Hence in the manuscript we settled with a rate of $\simeq 1\%$ FDR at the gene level and $\simeq 5\%$ FDR at the site level.

### 6 Rate of adaptation enrichment with polyDFE

The probability to sample allele at a given frequency (before fixation or extinction) is informative of its scaled selection coefficient at the population scale ($\beta$). Pooled across many sites, the SFS is thus informative on the underlying $\beta$ of mutations, given we have a neutral expectation. In this configuration, a single $\beta$ for all sampled mutations is biologically not realistic. Accordingly, a distribution of fitness effects of mutations (DFE) is assumed, usually modeled as a continuous distribution[1, 2]. In this study, we used the software polyDFE[3, 4] with model C (continuous distribution) and D (discrete distribution).

PolyDFE requires one SFS for non-synonymous mutations and one for synonymous mutations (neutral expectation), as well as the total number of sites on which each SFS has been sampled. From the SFS and the number of sites for both synonymous and non-synonymous changes, polyDFE estimates parameters of the DFE ($\phi(\beta)$) using maximum likelihood. The estimated DFE allows to subsequently computed the rate of adaptive evolution $\omega_{\mathrm{A}}^{\mathrm{polyDFE}}$.

##### polyDFE Model C

In model C, the DFE ($\phi$) is given by a mixture of a $\Gamma$ and Exponential distributions, parameterized by $\beta_d$ , $b$, $p_b$ and $\beta_b$ as:

$$\phi(\beta) = \begin{cases} (1-p_b)\, f_\Gamma(-\beta; -\beta_d, b) & \text{if } \beta \leq 0, \\ p_b f_e(\beta; \beta_b) & \text{if } \beta > 0, \end{cases} \tag{3}$$

where $\beta_d < 0$ is the estimated mean of the DFE for $\beta \leq 0$, $b > 0$ is the shape of the $\Gamma$ distribution fixed to 1.0, $0 \leq p_b \leq 1$ is the estimated probability that $\beta > 0$, $\beta_b > 0$ is the mean of the DFE for $\beta > 0$ fixed to 4.0, and $f_\Gamma(x; m, b)$ is the density of the $\Gamma$ distribution with mean m and shape b, while $f_e(x; m)$ is the density of the Exponential distribution with mean $m$.

##### polyDFE Model D

In model D, the DFE ($\phi$) is given as a discrete distribution, where the selection coefficients can take one of $\beta_i$ distinct values, $1 \leq i \leq K$, where each value $\beta_i$ has probability $p_i$, with

$$\sum_{i=0}^{K} p_i = 1. \tag{4}$$

#### 6.1 PolyDFE model C - including divergence data

The rate of adaptation $\omega_{\mathrm{A}}^{\mathrm{polyDFE}}$ is computed as the difference between the total rate of evolution $\omega = d_N/d_S$ obtained from divergence data and the rate of non-adaptive evolution ($\omega_{\mathrm{NA}}(\phi)$) obtained from polymorphism data as in Tataru *et al.* [3]:

$$\omega_{\mathrm{A}}^{\mathrm{polyDFE}} = \omega - \omega_{\mathrm{NA}}(\phi), \tag{5}$$

$$= d_N/d_S - \omega_{\mathrm{NA}}(\phi). \tag{6}$$

Formally, $\omega_{\mathrm{NA}}(\phi)$ is computed as the average fixation probability of mutations ($\mathbb{P}_{\mathrm{fix}}(\beta)$) over the probability distribution given by the DFE $\phi(\beta)$, taken only for the negatively selected mutations ($\beta < 0$) as:

$$\omega_{\mathrm{NA}}(\phi) = \int_{-\infty}^{0} \mathbb{P}_{\mathrm{fix}}(\beta)\phi(\beta)\mathrm{d}\beta, \tag{7}$$

$$= \int_{-\infty}^{0} \frac{\beta}{1-\mathrm{e}^{-\beta}}\phi(\beta)\mathrm{d}\beta, \tag{8}$$

$$= \int_{-\infty}^{0} \frac{\beta}{1-\mathrm{e}^{-\beta}}\,(1-p_b)\, f_\Gamma(-\beta; -\beta_d, b)\mathrm{d}\beta. \tag{9}$$

(10)

Altogether, $\omega_{\mathrm{A}}^{\mathrm{polyDFE}}$ is given as:

$$\omega_{\mathrm{A}}^{\mathrm{polyDFE}} = d_N/d_S - \omega_{\mathrm{NA}}(\phi), \tag{11}$$

$$= d_N/d_S - \int_{-\infty}^{0} \frac{\beta}{1-\mathrm{e}^{-\beta}} \phi(\beta)\mathrm{d}\beta. \tag{12}$$

##### Mutation-selection codon model at gene level ($\alpha$=0.025)

Figure S10:

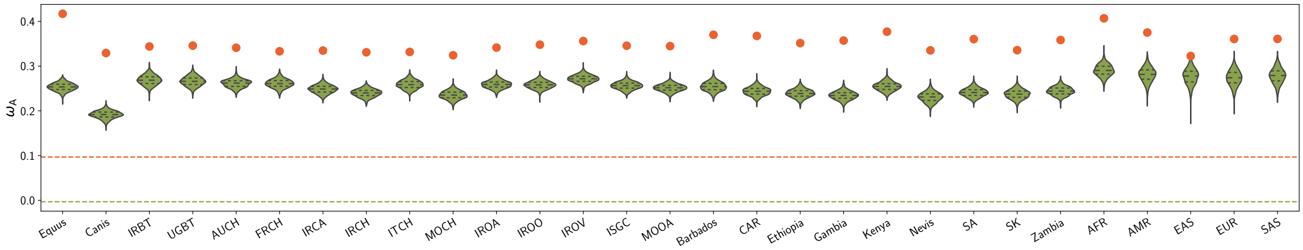


Table S11:

| Population | Species | $\omega_{\mathrm{A}}$ Adaptive | $\langle\omega_{\mathrm{A}}\rangle$ Nearly-neutral | $\Delta\omega_{\mathrm{A}}$ | $p_{\mathrm{v}}$ | $p_{\mathrm{v}}^{\mathrm{adj}}$ | $\frac{\Delta\omega_{\mathrm{A}}}{\omega_{\mathrm{A}}^{\mathrm{phy}}}$ | $\pi_{\mathrm{S}}$ |
|---|---|---|---|---|---|---|---|---|
| Diverse (Equus) | Equus caballus | 0.417 | 0.253 | 0.163 | 0.0 | **0.0*** | 1.702 | 0.00093 |
| Diverse (Canis) | Canis familiaris | 0.329 | 0.191 | 0.138 | 0.0 | **0.0*** | 1.335 | 0.001 |
| Iran (IRBT) | Bos taurus | 0.344 | 0.269 | 0.075 | 0.0 | **0.0*** | 0.748 | 0.003 |
| Uganda (UGBT) | Bos taurus | 0.346 | 0.266 | 0.080 | 0.0 | **0.0*** | 0.784 | 0.003 |
| Australia (AUCH) | Capra hircus | 0.341 | 0.261 | 0.080 | 0.0 | **0.0*** | 0.820 | 0.00099 |
| France (FRCH) | Capra hircus | 0.333 | 0.261 | 0.072 | 0.0 | **0.0*** | 0.744 | 0.00097 |
| Iran (IRCA) | Capra aegagrus | 0.335 | 0.248 | 0.086 | 0.0 | **0.0*** | 0.892 | 0.001 |
| Iran (IRCH) | Capra hircus | 0.331 | 0.240 | 0.091 | 0.0 | **0.0*** | 0.921 | 0.001 |
| Italy (ITCH) | Capra hircus | 0.332 | 0.259 | 0.073 | 0.0 | **0.0*** | 0.762 | 0.001 |
| Morocco (MOCH) | Capra hircus | 0.324 | 0.235 | 0.089 | 0.0 | **0.0*** | 0.886 | 0.001 |
| Iran (IROA) | Ovis aries | 0.341 | 0.259 | 0.082 | 0.0 | **0.0*** | 0.823 | 0.002 |
| Iran (IROO) | Ovis orientalis | 0.348 | 0.258 | 0.090 | 0.0 | **0.0*** | 0.901 | 0.003 |
| Iran (IROV) | Ovis vignei | 0.356 | 0.272 | 0.084 | 0.0 | **0.0*** | 0.860 | 0.002 |
| Various (ISGC) | Ovis aries | 0.346 | 0.256 | 0.089 | 0.0 | **0.0*** | 0.888 | 0.003 |
| Morocco (MOOA) | Ovis aries | 0.345 | 0.252 | 0.093 | 0.0 | **0.0*** | 0.921 | 0.002 |
| Barbados | Chlorocebus sabaeus | 0.370 | 0.254 | 0.116 | 0.0 | **0.0*** | 1.156 | 0.001 |
| Central African Republic (CAR) | Chlorocebus sabaeus | 0.367 | 0.244 | 0.124 | 0.0 | **0.0*** | 1.234 | 0.002 |
| Ethiopia | Chlorocebus sabaeus | 0.351 | 0.239 | 0.112 | 0.0 | **0.0*** | 1.126 | 0.002 |
| Gambia | Chlorocebus sabaeus | 0.357 | 0.234 | 0.123 | 0.0 | **0.0*** | 1.219 | 0.002 |
| Kenya | Chlorocebus sabaeus | 0.377 | 0.255 | 0.122 | 0.0 | **0.0*** | 1.245 | 0.001 |
| Nevis | Chlorocebus sabaeus | 0.335 | 0.231 | 0.104 | 0.0 | **0.0*** | 1.059 | 0.001 |
| South Africa (SA) | Chlorocebus sabaeus | 0.360 | 0.241 | 0.119 | 0.0 | **0.0*** | 1.184 | 0.002 |
| Saint Kitts (SK) | Chlorocebus sabaeus | 0.336 | 0.237 | 0.098 | 0.0 | **0.0*** | 0.983 | 0.001 |
| Zambia | Chlorocebus sabaeus | 0.358 | 0.244 | 0.114 | 0.0 | **0.0*** | 1.137 | 0.002 |
| African (AFR) | Homo sapiens | 0.407 | 0.290 | 0.116 | 0.0 | **0.0*** | 1.131 | 0.00071 |
| Ad Mixed American (AMR) | Homo sapiens | 0.375 | 0.281 | 0.094 | 0.0 | **0.0*** | 0.917 | 0.00056 |
| East Asian (EAS) | Homo sapiens | 0.323 | 0.276 | 0.046 | 0.0 | **0.0*** | 0.449 | 0.00051 |
| European (EUR) | Homo sapiens | 0.360 | 0.273 | 0.087 | 0.0 | **0.0*** | 0.847 | 0.00054 |
| South Asian (SAS) | Homo sapiens | 0.361 | 0.278 | 0.082 | 0.0 | **0.0*** | 0.805 | 0.00056 |

##### Mutation-selection codon model at site level ($\alpha$=0.025)

Figure S11:

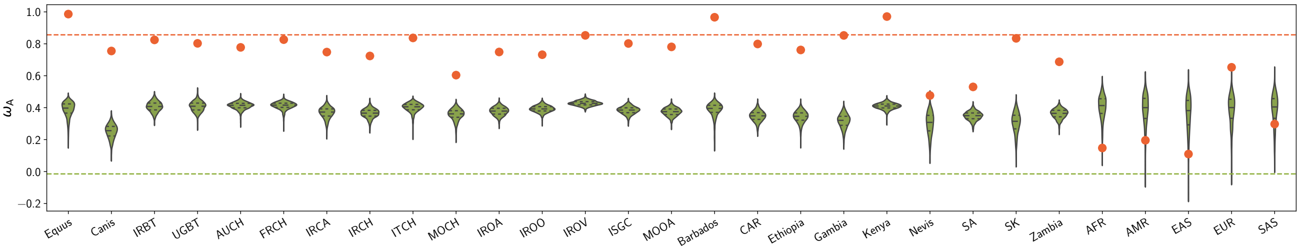


Table S12:

| Population | Species | $\omega_A$ Adaptive | $\langle\omega_A\rangle$ Nearly-neutral | $\Delta\omega_A$ | $p_v$ | $p_v^{adj}$ | $\frac{\Delta\omega_A}{\omega_A^{phy}}$ | $\pi_S$ |
|---|---|---|---|---|---|---|---|---|
| Diverse (Equus) | Equus caballus | 0.986 | 0.388 | 0.598 | 0.0 | **0.0*** | 0.688 | 0.00093 |
| Diverse (Canis) | Canis familiaris | 0.755 | 0.251 | 0.504 | 0.0 | **0.0*** | 0.575 | 0.001 |
| Iran (IRBT) | Bos taurus | 0.824 | 0.407 | 0.418 | 0.0 | **0.0*** | 0.480 | 0.003 |
| Uganda (UGBT) | Bos taurus | 0.803 | 0.407 | 0.396 | 0.0 | **0.0*** | 0.455 | 0.003 |
| Australia (AUCH) | Capra hircus | 0.778 | 0.412 | 0.366 | 0.0 | **0.0*** | 0.417 | 0.00099 |
| France (FRCH) | Capra hircus | 0.826 | 0.413 | 0.413 | 0.0 | **0.0*** | 0.472 | 0.00097 |
| Iran (IRCA) | Capra aegagrus | 0.748 | 0.368 | 0.380 | 0.0 | **0.0*** | 0.436 | 0.001 |
| Iran (IRCH) | Capra hircus | 0.724 | 0.364 | 0.360 | 0.0 | **0.0*** | 0.412 | 0.001 |
| Italy (ITCH) | Capra hircus | 0.837 | 0.401 | 0.436 | 0.0 | **0.0*** | 0.500 | 0.001 |
| Morocco (MOCH) | Capra hircus | 0.604 | 0.357 | 0.247 | 0.0 | **0.0*** | 0.282 | 0.001 |
| Iran (IROA) | Ovis aries | 0.749 | 0.378 | 0.371 | 0.0 | **0.0*** | 0.425 | 0.002 |
| Iran (IROO) | Ovis orientalis | 0.732 | 0.392 | 0.339 | 0.0 | **0.0*** | 0.390 | 0.003 |
| Iran (IROV) | Ovis vignei | 0.853 | 0.428 | 0.425 | 0.0 | **0.0*** | 0.489 | 0.002 |
| Various (ISGC) | Ovis aries | 0.802 | 0.384 | 0.418 | 0.0 | **0.0*** | 0.478 | 0.003 |
| Morocco (MOOA) | Ovis aries | 0.781 | 0.373 | 0.407 | 0.0 | **0.0*** | 0.466 | 0.002 |
| Barbados | Chlorocebus sabaeus | 0.966 | 0.386 | 0.580 | 0.0 | **0.0*** | 0.668 | 0.001 |
| Central African Republic (CAR) | Chlorocebus sabaeus | 0.799 | 0.347 | 0.452 | 0.0 | **0.0*** | 0.521 | 0.002 |
| Ethiopia | Chlorocebus sabaeus | 0.762 | 0.343 | 0.419 | 0.0 | **0.0*** | 0.482 | 0.002 |
| Gambia | Chlorocebus sabaeus | 0.852 | 0.316 | 0.536 | 0.0 | **0.0*** | 0.618 | 0.002 |
| Kenya | Chlorocebus sabaeus | 0.971 | 0.410 | 0.561 | 0.0 | **0.0*** | 0.649 | 0.001 |
| Nevis | Chlorocebus sabaeus | 0.476 | 0.300 | 0.176 | 0.0 | **0.0*** | 0.203 | 0.001 |
| South Africa (SA) | Chlorocebus sabaeus | 0.530 | 0.348 | 0.182 | 0.0 | **0.0*** | 0.209 | 0.002 |
| Saint Kitts (SK) | Chlorocebus sabaeus | 0.834 | 0.307 | 0.527 | 0.0 | **0.0*** | 0.607 | 0.001 |
| Zambia | Chlorocebus sabaeus | 0.687 | 0.362 | 0.326 | 0.0 | **0.0*** | 0.375 | 0.002 |
| African (AFR) | Homo sapiens | 0.148 | 0.404 | −0.256 | 0.995 | 1.000 | −0.294 | 0.00071 |
| Ad Mixed American (AMR) | Homo sapiens | 0.196 | 0.388 | −0.192 | 0.961 | 1.000 | −0.220 | 0.00056 |
| East Asian (EAS) | Homo sapiens | 0.110 | 0.360 | −0.250 | 0.977 | 1.000 | −0.287 | 0.00051 |
| European (EUR) | Homo sapiens | 0.653 | 0.385 | 0.268 | 0.0 | **0.0*** | 0.307 | 0.00054 |
| South Asian (SAS) | Homo sapiens | 0.298 | 0.388 | −0.090 | 0.840 | 1.000 | −0.104 | 0.00056 |

- At the gene level, sets of genes supposedly under a nearly-neutral regime have an average value of $\omega_A^{\text{polyDFE}}$ (average across the replicates) in the range $[0.19, 0.29]$ across the populations, while $\omega_A^{\text{polyDFE}}$ for genes under adaptation (at the phylogenetic scale) is in the range $[0.32, 0.42]$ across the populations.

- At the site level, sets of sites supposedly under a nearly-neutral regime have an average value of $\omega_A^{\text{polyDFE}}$ (average across the replicates) in the range $[0.25, 0.42]$ across the populations, while $\omega_A^{\text{polyDFE}}$ for genes under adaptation (at the phylogenetic scale) is in the range $[0.11, 0.99]$ across the populations.

$\omega_A^{\text{polyDFE}}$ computed with polyDFE is higher than $\omega_A$ computed as the McDonald & Kreitman [5] statistic (figure 3 and table 1 in the main manuscript), suggesting that polyDFE has higher sensitivity to detect adaptation at the population-genetic scale. However, the genes and sites supposedly under a nearly-neutral regime have all values of $\omega_A^{\text{polyDFE}}$ greater than 0, suggesting that the higher sensitivity (true positive rate) also results in a lower specificity (true negative rate). Altogether, the statistical test for the enrichment of $\omega_A$ between the set of adaptive and nearly-neutral genes and sites gives similar results whether computed by polyDFE ($\omega_A^{\text{polyDFE}}$) or as McDonald & Kreitman [5] statistic.

#### 6.2 PolyDFE model C - polymorphism data alone with $\beta > 0$

As in Tataru *et al.* [3], the rate of adaptation $\omega_{\mathrm{A}}^{\mathrm{polyDFE}}$ can also be estimated from the polymorphism data alone, computed as the average fixation probability $\mathbb{P}_{\mathrm{fix}}(\beta)$ over the probability given the DFE ($\phi(\beta)$), only for the positively selected mutations ($\beta > 0$), as:

$$\omega_{\mathrm{A}}^{\mathrm{polyDFE}} = \int_0^{+\infty} \mathbb{P}_{\mathrm{fix}}(\beta)\phi(\beta)\mathrm{d}\beta, \tag{13}$$

$$= \int_0^{+\infty} \frac{\beta}{1 - \mathrm{e}^{-\beta}}\phi(\beta)\mathrm{d}\beta, \tag{14}$$

$$= \int_0^{+\infty} \frac{\beta}{1 - \mathrm{e}^{-\beta}} p_b f_e(\beta; \beta_b)\mathrm{d}\beta. \tag{15}$$

**Mutation-selection codon model at gene level ($\alpha$=0.025)**

Figure S12:

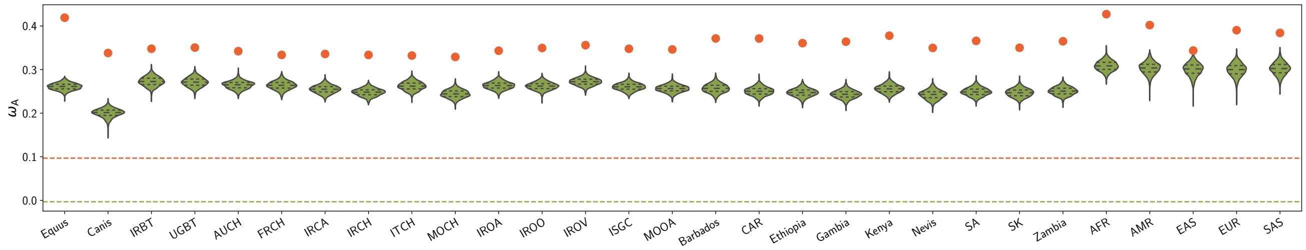


Table S13:

| Population | Species | $\omega_{\mathrm{A}}$ Adaptive | $\langle\omega_{\mathrm{A}}\rangle$ Nearly-neutral | $\Delta\omega_{\mathrm{A}}$ | $p_{\mathrm{v}}$ | $p_{\mathrm{v}}^{\mathrm{adj}}$ | $\frac{\Delta\omega_{\mathrm{A}}}{\omega_{\mathrm{A}}^{\mathrm{phy}}}$ | $\pi_{\mathrm{S}}$ |
|---|---|---|---|---|---|---|---|---|
| Diverse (Equus) | Equus caballus | 0.419 | 0.261 | 0.158 | 0.0 | **0.0*** | 1.645 | 0.00093 |
| Diverse (Canis) | Canis familiaris | 0.338 | 0.201 | 0.138 | 0.0 | **0.0*** | 1.333 | 0.001 |
| Iran (IRBT) | Bos taurus | 0.348 | 0.273 | 0.075 | 0.0 | **0.0*** | 0.744 | 0.003 |
| Uganda (UGBT) | Bos taurus | 0.351 | 0.271 | 0.079 | 0.0 | **0.0*** | 0.779 | 0.003 |
| Australia (AUCH) | Capra hircus | 0.342 | 0.265 | 0.078 | 0.0 | **0.0*** | 0.799 | 0.00099 |
| France (FRCH) | Capra hircus | 0.334 | 0.264 | 0.070 | 0.0 | **0.0*** | 0.721 | 0.00097 |
| Iran (IRCA) | Capra aegagrus | 0.336 | 0.255 | 0.081 | 0.0 | **0.0*** | 0.837 | 0.001 |
| Iran (IRCH) | Capra hircus | 0.334 | 0.248 | 0.086 | 0.0 | **0.0*** | 0.866 | 0.001 |
| Italy (ITCH) | Capra hircus | 0.332 | 0.262 | 0.070 | 0.0 | **0.0*** | 0.730 | 0.001 |
| Morocco (MOCH) | Capra hircus | 0.329 | 0.244 | 0.085 | 0.0 | **0.0*** | 0.844 | 0.001 |
| Iran (IROA) | Ovis aries | 0.343 | 0.263 | 0.080 | 0.0 | **0.0*** | 0.799 | 0.002 |
| Iran (IROO) | Ovis orientalis | 0.350 | 0.262 | 0.087 | 0.0 | **0.0*** | 0.877 | 0.003 |
| Iran (IROV) | Ovis vignei | 0.356 | 0.272 | 0.084 | 0.0 | **0.0*** | 0.853 | 0.002 |
| Various (ISGC) | Ovis aries | 0.348 | 0.260 | 0.087 | 0.0 | **0.0*** | 0.868 | 0.003 |
| Morocco (MOOA) | Ovis aries | 0.346 | 0.256 | 0.090 | 0.0 | **0.0*** | 0.888 | 0.002 |
| Barbados | Chlorocebus sabaeus | 0.371 | 0.257 | 0.114 | 0.0 | **0.0*** | 1.145 | 0.001 |
| Central African Republic (CAR) | Chlorocebus sabaeus | 0.371 | 0.250 | 0.121 | 0.0 | **0.0*** | 1.208 | 0.002 |
| Ethiopia | Chlorocebus sabaeus | 0.361 | 0.247 | 0.114 | 0.0 | **0.0*** | 1.137 | 0.002 |
| Gambia | Chlorocebus sabaeus | 0.364 | 0.243 | 0.121 | 0.0 | **0.0*** | 1.199 | 0.002 |
| Kenya | Chlorocebus sabaeus | 0.378 | 0.256 | 0.122 | 0.0 | **0.0*** | 1.240 | 0.001 |
| Nevis | Chlorocebus sabaeus | 0.350 | 0.242 | 0.107 | 0.0 | **0.0*** | 1.087 | 0.001 |
| South Africa (SA) | Chlorocebus sabaeus | 0.366 | 0.249 | 0.117 | 0.0 | **0.0*** | 1.160 | 0.002 |
| Saint Kitts (SK) | Chlorocebus sabaeus | 0.350 | 0.247 | 0.103 | 0.0 | **0.0*** | 1.033 | 0.001 |
| Zambia | Chlorocebus sabaeus | 0.365 | 0.251 | 0.114 | 0.0 | **0.0*** | 1.139 | 0.002 |
| African (AFR) | Homo sapiens | 0.427 | 0.309 | 0.119 | 0.0 | **0.0*** | 1.154 | 0.00071 |
| Ad Mixed American (AMR) | Homo sapiens | 0.402 | 0.303 | 0.099 | 0.0 | **0.0*** | 0.963 | 0.00056 |
| East Asian (EAS) | Homo sapiens | 0.344 | 0.300 | 0.043 | 0.0 | **0.0*** | 0.421 | 0.00051 |
| European (EUR) | Homo sapiens | 0.390 | 0.299 | 0.091 | 0.0 | **0.0*** | 0.886 | 0.00054 |
| South Asian (SAS) | Homo sapiens | 0.384 | 0.302 | 0.082 | 0.0 | **0.0*** | 0.799 | 0.00056 |

##### Mutation-selection codon model at site level ($\alpha$=0.025)

Figure S13:

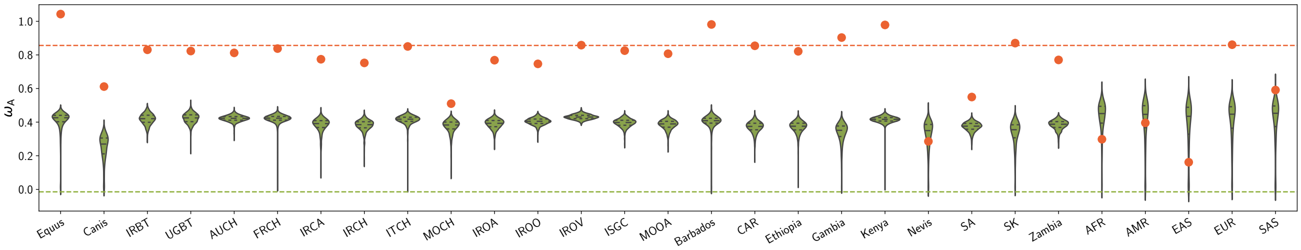


Table S14:

| Population | Species | $\omega_{\text{A}}$ Adaptive | $\langle\omega_{\text{A}}\rangle$ Nearly-neutral | $\Delta\omega_{\text{A}}$ | $p_{\text{v}}$ | $p_{\text{v}}^{\text{adj}}$ | $\frac{\Delta\omega_{\text{A}}}{\omega_{\text{A}}^{\text{phy}}}$ | $\pi_{\text{S}}$ |
|---|---|---|---|---|---|---|---|---|
| Diverse (Equus) | Equus caballus | 1.043 | 0.407 | 0.636 | 0.0 | **0.0*** | 0.732 | 0.00093 |
| Diverse (Canis) | Canis familiaris | 0.612 | 0.249 | 0.363 | 0.0 | **0.0*** | 0.414 | 0.001 |
| Iran (IRBT) | Bos taurus | 0.830 | 0.418 | 0.412 | 0.0 | **0.0*** | 0.474 | 0.003 |
| Uganda (UGBT) | Bos taurus | 0.823 | 0.423 | 0.400 | 0.0 | **0.0*** | 0.461 | 0.003 |
| Australia (AUCH) | Capra hircus | 0.812 | 0.422 | 0.390 | 0.0 | **0.0*** | 0.446 | 0.00099 |
| France (FRCH) | Capra hircus | 0.838 | 0.419 | 0.418 | 0.0 | **0.0*** | 0.478 | 0.00097 |
| Iran (IRCA) | Capra aegagrus | 0.774 | 0.382 | 0.392 | 0.0 | **0.0*** | 0.450 | 0.001 |
| Iran (IRCH) | Capra hircus | 0.752 | 0.381 | 0.372 | 0.0 | **0.0*** | 0.425 | 0.001 |
| Italy (ITCH) | Capra hircus | 0.850 | 0.412 | 0.438 | 0.0 | **0.0*** | 0.502 | 0.001 |
| Morocco (MOCH) | Capra hircus | 0.510 | 0.374 | 0.136 | 0.0 | **0.0*** | 0.156 | 0.001 |
| Iran (IROA) | Ovis aries | 0.769 | 0.390 | 0.379 | 0.0 | **0.0*** | 0.434 | 0.002 |
| Iran (IROO) | Ovis orientalis | 0.747 | 0.404 | 0.343 | 0.0 | **0.0*** | 0.394 | 0.003 |
| Iran (IROV) | Ovis vignei | 0.858 | 0.431 | 0.427 | 0.0 | **0.0*** | 0.492 | 0.002 |
| Various (ISGC) | Ovis aries | 0.825 | 0.395 | 0.430 | 0.0 | **0.0*** | 0.492 | 0.003 |
| Morocco (MOOA) | Ovis aries | 0.807 | 0.385 | 0.422 | 0.0 | **0.0*** | 0.483 | 0.002 |
| Barbados | Chlorocebus sabaeus | 0.981 | 0.399 | 0.582 | 0.0 | **0.0*** | 0.670 | 0.001 |
| Central African Republic (CAR) | Chlorocebus sabaeus | 0.854 | 0.370 | 0.484 | 0.0 | **0.0*** | 0.558 | 0.002 |
| Ethiopia | Chlorocebus sabaeus | 0.821 | 0.370 | 0.452 | 0.0 | **0.0*** | 0.520 | 0.002 |
| Gambia | Chlorocebus sabaeus | 0.903 | 0.338 | 0.565 | 0.0 | **0.0*** | 0.651 | 0.002 |
| Kenya | Chlorocebus sabaeus | 0.978 | 0.416 | 0.562 | 0.0 | **0.0*** | 0.650 | 0.001 |
| Nevis | Chlorocebus sabaeus | 0.286 | 0.326 | −0.040 | 0.771 | 1.000 | −0.047 | 0.001 |
| South Africa (SA) | Chlorocebus sabaeus | 0.549 | 0.373 | 0.177 | 0.0 | **0.0*** | 0.203 | 0.002 |
| Saint Kitts (SK) | Chlorocebus sabaeus | 0.870 | 0.332 | 0.538 | 0.0 | **0.0*** | 0.620 | 0.001 |
| Zambia | Chlorocebus sabaeus | 0.770 | 0.384 | 0.386 | 0.0 | **0.0*** | 0.445 | 0.002 |
| African (AFR) | Homo sapiens | 0.298 | 0.425 | −0.127 | 0.898 | 1.000 | −0.146 | 0.00071 |
| Ad Mixed American (AMR) | Homo sapiens | 0.396 | 0.406 | −0.010 | 0.681 | 1.000 | −0.011 | 0.00056 |
| East Asian (EAS) | Homo sapiens | 0.161 | 0.382 | −0.221 | 0.882 | 1.000 | −0.253 | 0.00051 |
| European (EUR) | Homo sapiens | 0.861 | 0.407 | 0.454 | 0.0 | **0.0*** | 0.521 | 0.00054 |
| South Asian (SAS) | Homo sapiens | 0.591 | 0.409 | 0.182 | 0.001 | **0.005*** | 0.208 | 0.00056 |

- At the gene level, sets of genes supposedly under a nearly-neutral regime have an average value of $\omega_{\text{A}}^{\text{polyDFE}}$ (average across the replicates) in the range $[0.20, 0.31]$ across the populations, while $\omega_{\text{A}}^{\text{polyDFE}}$ for genes under adaptation (at the phylogenetic scale) is in the range $[0.32, 0.43]$ across the populations.
- At the site level, sets of sites supposedly under a nearly-neutral regime have an average value of $\omega_{\text{A}}^{\text{polyDFE}}$ (average across the replicates) in the range $[0.24, 0.44]$ across the populations, while $\omega_{\text{A}}^{\text{polyDFE}}$ for genes under adaptation (at the phylogenetic scale) is in the range $[0.16, 1.04]$ across the populations.

The estimation of $\omega_{\text{A}}^{\text{polyDFE}}$ computed with polyDFE using polymorphism data alone is quite consistent with the estimation combining polymorphism and divergence (previous section). However, we higher variance in the estimation (violin plots are more extended for the set of nearly-neutral replicates, particularly noticeable for sites), reducing the statistical power for the enrichment test of $\omega_{\text{A}}^{\text{polyDFE}}$ between the set of adaptive and nearly-neutral genes and sites.

#### 6.3 PolyDFE model C - polymorphism data alone with $\beta > 5$

The definition for a positively selected mutation by the criterion $\beta > 0$ is also open to interpretation, and the integration limit can be set to a strictly positive value (e.g. 1, 3 or 5) instead of 0[3, 6]. The reasoning is that mutations with a positive selection coefficient that is not very large are not necessarily advantageous mutations. In Galtier [6], the threshold $\beta > 5$ is used and the rate of adaptation is thus:

$$\omega_{\mathrm{A}}^{\mathrm{polyDFE}} = \int_{5}^{+\infty} \frac{\beta}{1 - \mathrm{e}^{-\beta}} p_b f_e(\beta; \beta_b) \mathrm{d}\beta. \tag{16}$$

##### Mutation-selection codon model at gene level ($\alpha$=0.025)

Figure S14:

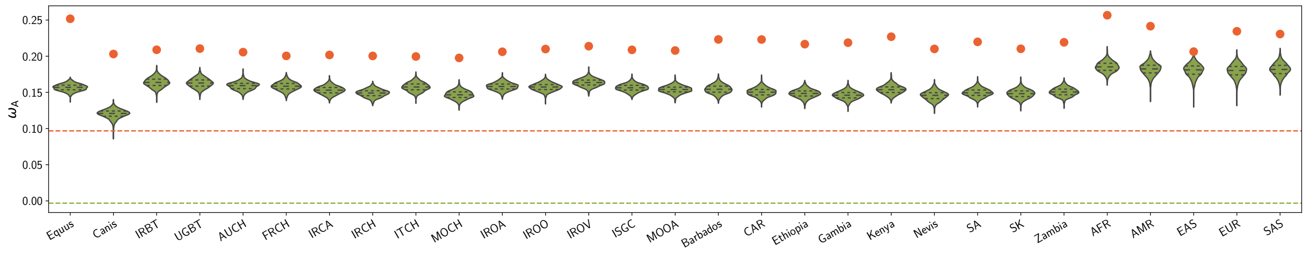


Table S15:

| Population | Species | $\omega_{\mathrm{A}}$ Adaptive | $\langle\omega_{\mathrm{A}}\rangle$ Nearly-neutral | $\Delta\omega_{\mathrm{A}}$ | $p_{\mathrm{v}}$ | $p_{\mathrm{v}}^{\mathrm{adj}}$ | $\frac{\Delta\omega_{\mathrm{A}}}{\omega_{\mathrm{A}}^{\mathrm{phy}}}$ | $\pi_{\mathrm{S}}$ |
|---|---|---|---|---|---|---|---|---|
| Diverse (Equus) | Equus caballus | 0.252 | 0.157 | 0.095 | 0.0 | **0.0*** | 0.989 | 0.00093 |
| Diverse (Canis) | Canis familiaris | 0.203 | 0.120 | 0.083 | 0.0 | **0.0*** | 0.802 | 0.001 |
| Iran (IRBT) | Bos taurus | 0.209 | 0.164 | 0.045 | 0.0 | **0.0*** | 0.447 | 0.003 |
| Uganda (UGBT) | Bos taurus | 0.211 | 0.163 | 0.048 | 0.0 | **0.0*** | 0.468 | 0.003 |
| Australia (AUCH) | Capra hircus | 0.206 | 0.159 | 0.047 | 0.0 | **0.0*** | 0.480 | 0.00099 |
| France (FRCH) | Capra hircus | 0.200 | 0.158 | 0.042 | 0.0 | **0.0*** | 0.433 | 0.00097 |
| Iran (IRCA) | Capra aegagrus | 0.202 | 0.153 | 0.049 | 0.0 | **0.0*** | 0.503 | 0.001 |
| Iran (IRCH) | Capra hircus | 0.200 | 0.149 | 0.051 | 0.0 | **0.0*** | 0.521 | 0.001 |
| Italy (ITCH) | Capra hircus | 0.200 | 0.158 | 0.042 | 0.0 | **0.0*** | 0.439 | 0.001 |
| Morocco (MOCH) | Capra hircus | 0.198 | 0.147 | 0.051 | 0.0 | **0.0*** | 0.507 | 0.001 |
| Iran (IROA) | Ovis aries | 0.206 | 0.158 | 0.048 | 0.0 | **0.0*** | 0.480 | 0.002 |
| Iran (IROO) | Ovis orientalis | 0.210 | 0.157 | 0.053 | 0.0 | **0.0*** | 0.527 | 0.003 |
| Iran (IROV) | Ovis vignei | 0.214 | 0.164 | 0.050 | 0.0 | **0.0*** | 0.513 | 0.002 |
| Various (ISGC) | Ovis aries | 0.209 | 0.156 | 0.052 | 0.0 | **0.0*** | 0.522 | 0.003 |
| Morocco (MOOA) | Ovis aries | 0.208 | 0.154 | 0.054 | 0.0 | **0.0*** | 0.534 | 0.002 |
| Barbados | Chlorocebus sabaeus | 0.223 | 0.154 | 0.069 | 0.0 | **0.0*** | 0.688 | 0.001 |
| Central African Republic (CAR) | Chlorocebus sabaeus | 0.223 | 0.150 | 0.073 | 0.0 | **0.0*** | 0.726 | 0.002 |
| Ethiopia | Chlorocebus sabaeus | 0.217 | 0.148 | 0.068 | 0.0 | **0.0*** | 0.683 | 0.002 |
| Gambia | Chlorocebus sabaeus | 0.219 | 0.146 | 0.073 | 0.0 | **0.0*** | 0.721 | 0.002 |
| Kenya | Chlorocebus sabaeus | 0.227 | 0.154 | 0.073 | 0.0 | **0.0*** | 0.745 | 0.001 |
| Nevis | Chlorocebus sabaeus | 0.210 | 0.146 | 0.065 | 0.0 | **0.0*** | 0.654 | 0.001 |
| South Africa (SA) | Chlorocebus sabaeus | 0.220 | 0.150 | 0.070 | 0.0 | **0.0*** | 0.698 | 0.002 |
| Saint Kitts (SK) | Chlorocebus sabaeus | 0.210 | 0.148 | 0.062 | 0.0 | **0.0*** | 0.621 | 0.001 |
| Zambia | Chlorocebus sabaeus | 0.219 | 0.151 | 0.069 | 0.0 | **0.0*** | 0.684 | 0.002 |
| African (AFR) | Homo sapiens | 0.257 | 0.185 | 0.071 | 0.0 | **0.0*** | 0.693 | 0.00071 |
| Ad Mixed American (AMR) | Homo sapiens | 0.242 | 0.182 | 0.059 | 0.0 | **0.0*** | 0.578 | 0.00056 |
| East Asian (EAS) | Homo sapiens | 0.206 | 0.180 | 0.026 | 0.0 | **0.0*** | 0.252 | 0.00051 |
| European (EUR) | Homo sapiens | 0.234 | 0.180 | 0.055 | 0.0 | **0.0*** | 0.532 | 0.00054 |
| South Asian (SAS) | Homo sapiens | 0.231 | 0.182 | 0.049 | 0.0 | **0.0*** | 0.480 | 0.00056 |

##### Mutation-selection codon model at site level ($\alpha$=0.025)

Figure S15:

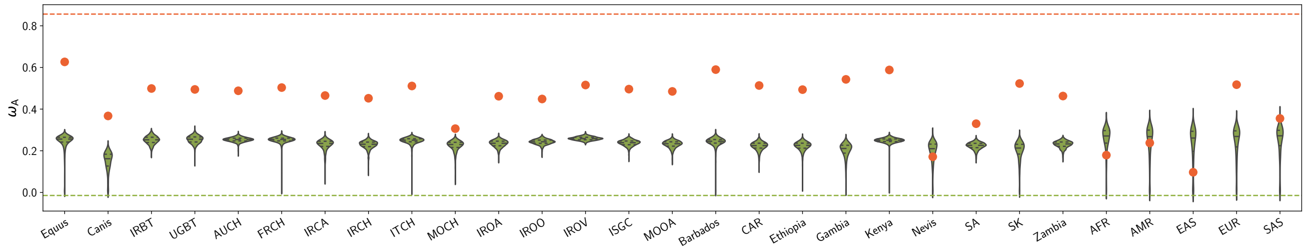


Table S16:

| Population | Species | $\omega_{\mathrm{A}}$ Adaptive | $\langle\omega_{\mathrm{A}}\rangle$ Nearly-neutral | $\Delta\omega_{\mathrm{A}}$ | $p_{\mathrm{v}}$ | $p_{\mathrm{v}}^{\mathrm{adj}}$ | $\frac{\Delta\omega_{\mathrm{A}}}{\omega_{\mathrm{A}}^{\mathrm{phy}}}$ | $\pi_{\mathrm{S}}$ |
|---|---|---|---|---|---|---|---|---|
| Diverse (Equus) | Equus caballus | 0.627 | 0.244 | 0.382 | 0.0 | **0.0*** | 0.440 | 0.00093 |
| Diverse (Canis) | Canis familiaris | 0.367 | 0.149 | 0.218 | 0.0 | **0.0*** | 0.249 | 0.001 |
| Iran (IRBT) | Bos taurus | 0.499 | 0.251 | 0.248 | 0.0 | **0.0*** | 0.285 | 0.003 |
| Uganda (UGBT) | Bos taurus | 0.495 | 0.254 | 0.241 | 0.0 | **0.0*** | 0.277 | 0.003 |
| Australia (AUCH) | Capra hircus | 0.488 | 0.253 | 0.235 | 0.0 | **0.0*** | 0.268 | 0.00099 |
| France (FRCH) | Capra hircus | 0.503 | 0.252 | 0.251 | 0.0 | **0.0*** | 0.287 | 0.00097 |
| Iran (IRCA) | Capra aegagrus | 0.465 | 0.229 | 0.236 | 0.0 | **0.0*** | 0.270 | 0.001 |
| Iran (IRCH) | Capra hircus | 0.452 | 0.229 | 0.223 | 0.0 | **0.0*** | 0.256 | 0.001 |
| Italy (ITCH) | Capra hircus | 0.511 | 0.248 | 0.263 | 0.0 | **0.0*** | 0.302 | 0.001 |
| Morocco (MOCH) | Capra hircus | 0.306 | 0.224 | 0.082 | 0.0 | **0.0*** | 0.094 | 0.001 |
| Iran (IROA) | Ovis aries | 0.462 | 0.234 | 0.228 | 0.0 | **0.0*** | 0.261 | 0.002 |
| Iran (IROO) | Ovis orientalis | 0.449 | 0.243 | 0.206 | 0.0 | **0.0*** | 0.237 | 0.003 |
| Iran (IROV) | Ovis vignei | 0.516 | 0.259 | 0.257 | 0.0 | **0.0*** | 0.296 | 0.002 |
| Various (ISGC) | Ovis aries | 0.496 | 0.237 | 0.259 | 0.0 | **0.0*** | 0.296 | 0.003 |
| Morocco (MOOA) | Ovis aries | 0.485 | 0.231 | 0.254 | 0.0 | **0.0*** | 0.290 | 0.002 |
| Barbados | Chlorocebus sabaeus | 0.590 | 0.240 | 0.350 | 0.0 | **0.0*** | 0.403 | 0.001 |
| Central African Republic (CAR) | Chlorocebus sabaeus | 0.513 | 0.222 | 0.291 | 0.0 | **0.0*** | 0.335 | 0.002 |
| Ethiopia | Chlorocebus sabaeus | 0.494 | 0.222 | 0.271 | 0.0 | **0.0*** | 0.313 | 0.002 |
| Gambia | Chlorocebus sabaeus | 0.543 | 0.203 | 0.340 | 0.0 | **0.0*** | 0.392 | 0.002 |
| Kenya | Chlorocebus sabaeus | 0.588 | 0.250 | 0.338 | 0.0 | **0.0*** | 0.391 | 0.001 |
| Nevis | Chlorocebus sabaeus | 0.171 | 0.196 | −0.024 | 0.773 | 1.000 | −0.028 | 0.001 |
| South Africa (SA) | Chlorocebus sabaeus | 0.330 | 0.224 | 0.106 | 0.0 | **0.0*** | 0.122 | 0.002 |
| Saint Kitts (SK) | Chlorocebus sabaeus | 0.523 | 0.199 | 0.323 | 0.0 | **0.0*** | 0.373 | 0.001 |
| Zambia | Chlorocebus sabaeus | 0.463 | 0.230 | 0.232 | 0.0 | **0.0*** | 0.267 | 0.002 |
| African (AFR) | Homo sapiens | 0.179 | 0.255 | −0.076 | 0.898 | 1.000 | −0.088 | 0.00071 |
| Ad Mixed American (AMR) | Homo sapiens | 0.238 | 0.244 | −0.006 | 0.681 | 1.000 | −0.007 | 0.00056 |
| East Asian (EAS) | Homo sapiens | 0.097 | 0.230 | −0.133 | 0.882 | 1.000 | −0.152 | 0.00051 |
| European (EUR) | Homo sapiens | 0.517 | 0.244 | 0.273 | 0.0 | **0.0*** | 0.313 | 0.00054 |
| South Asian (SAS) | Homo sapiens | 0.355 | 0.246 | 0.109 | 0.001 | **0.005*** | 0.125 | 0.00056 |

- At the gene level, sets of genes supposedly under a nearly-neutral regime have an average value of $\omega_{\mathrm{A}}^{\mathrm{polyDFE}}$ (average across the replicates) in the range $[0.12, 0.19]$ across the populations, while $\omega_{\mathrm{A}}^{\mathrm{polyDFE}}$ for genes under adaptation (at the phylogenetic scale) is in the range $[0.20, 0.26]$ across the populations.

- At the site level, sets of sites supposedly under a nearly-neutral regime have an average value of $\omega_{\mathrm{A}}^{\mathrm{polyDFE}}$ (average across the replicates) in the range $[0.14, 0.26]$ across the populations, while $\omega_{\mathrm{A}}^{\mathrm{polyDFE}}$ for genes under adaptation (at the phylogenetic scale) is in the range $[0.09, 0.63]$ across the populations.

The estimation of $\omega_{\mathrm{A}}^{\mathrm{polyDFE}}$ computed with polyDFE using polymorphism data alone and with a bound of $\beta > 5$ to consider a mutation as adaptive[6] is lower than $\omega_{\mathrm{A}}^{\mathrm{polyDFE}}$ computed with a bound of $\beta > 0$[3]. However, the genes and sites supposedly under a nearly-neutral regime still have values of $\omega_{\mathrm{A}}^{\mathrm{polyDFE}}$ greater than 0, suggesting that $\omega_{\mathrm{A}}^{\mathrm{polyDFE}}$ still has a lower specificity than the McDonald & Kreitman [5] statistic.

##### 6.4 PolyDFE model D - including divergence

Again, the rate of adaptation $\omega_{\mathrm{A}}^{\mathrm{polyDFE}}$ is computed as the difference between the total rate of evolution $\omega = d_N/d_S$ obtained from divergence data and the rate of non-adaptive evolution ($\omega_{\mathrm{NA}}(\phi)$) obtained from polymorphism data. Also, $\omega_{\mathrm{NA}}(\phi)$ is computed as the average fixation probability of mutations ($\mathbb{P}_{\mathrm{fix}}(\beta)$) over the probability distribution given by the DFE $\phi(\beta)$, taken only for the negatively selected mutations ($\beta < 0$) as:

$$\omega_{\mathrm{NA}}(\phi) = \sum_{i=1}^{K} \mathbb{1}_{]-\infty,0]}(\beta_i)\mathbb{P}_{\mathrm{fix}}(\beta_i)p_i, \tag{17}$$

$$= \sum_{i=1}^{K} \mathbb{1}_{]-\infty,0]}(\beta_i)\frac{\beta_i}{1-\mathrm{e}^{-\beta_i}}p_i, \tag{18}$$

where $\mathbb{1}_{]-\infty,0]}(\beta)$ is the indicator function:

$$\mathbb{1}_{]-\infty,0]}(\beta) = \begin{cases} 1 & \text{if } \beta \leq 0, \\ 0 & \text{if } \beta > 0. \end{cases} \tag{19}$$

Altogether, $\omega_{\mathrm{A}}^{\mathrm{polyDFE}}$ is given as:

$$\omega_{\mathrm{A}}^{\mathrm{polyDFE}} = d_N/d_S - \sum_{i=1}^{K} \mathbb{1}_{]-\infty,0]}(\beta_i)\frac{\beta_i}{1-\mathrm{e}^{-\beta_i}}p_i. \tag{20}$$

#### Mutation-selection codon model at gene level ($\alpha$=0.025)

Figure S16:

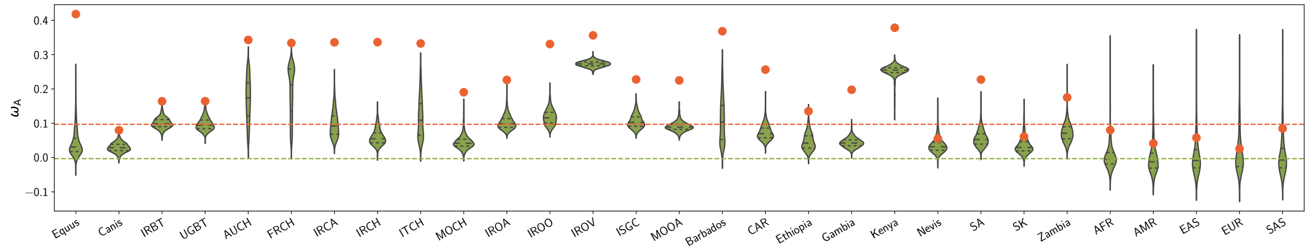


Table S17:

| Population | Species | $\omega_{\mathrm{A}}$ Adaptive | $\langle\omega_{\mathrm{A}}\rangle$ Nearly-neutral | $\Delta\omega_{\mathrm{A}}$ | $p_{\mathrm{v}}$ | $p_{\mathrm{v}}^{\mathrm{adj}}$ | $\frac{\Delta\omega_{\mathrm{A}}}{\omega_{\mathrm{A}}^{\mathrm{phy}}}$ | $\pi_{\mathrm{S}}$ |
|---|---|---|---|---|---|---|---|---|
| Diverse (Equus) | Equus caballus | 0.418 | 0.043 | 0.375 | 0.0 | **0.0*** | 3.907 | 0.00093 |
| Diverse (Canis) | Canis familiaris | 0.080 | 0.029 | 0.050 | 0.0 | **0.0*** | 0.488 | 0.001 |
| Iran (IRBT) | Bos taurus | 0.164 | 0.101 | 0.063 | 0.003 | **0.027*** | 0.627 | 0.003 |
| Uganda (UGBT) | Bos taurus | 0.164 | 0.098 | 0.066 | 0.001 | **0.011*** | 0.651 | 0.003 |
| Australia (AUCH) | Capra hircus | 0.343 | 0.170 | 0.173 | 0.0 | **0.0*** | 1.782 | 0.00099 |
| France (FRCH) | Capra hircus | 0.334 | 0.193 | 0.141 | 0.0 | **0.0*** | 1.455 | 0.00097 |
| Iran (IRCA) | Capra aegagrus | 0.336 | 0.097 | 0.238 | 0.0 | **0.0*** | 2.464 | 0.001 |
| Iran (IRCH) | Capra hircus | 0.336 | 0.060 | 0.277 | 0.0 | **0.0*** | 2.800 | 0.001 |
| Italy (ITCH) | Capra hircus | 0.332 | 0.116 | 0.216 | 0.0 | **0.0*** | 2.256 | 0.001 |
| Morocco (MOCH) | Capra hircus | 0.190 | 0.045 | 0.146 | 0.0 | **0.0*** | 1.453 | 0.001 |
| Iran (IROA) | Ovis aries | 0.226 | 0.102 | 0.124 | 0.0 | **0.0*** | 1.237 | 0.002 |
| Iran (IROO) | Ovis orientalis | 0.331 | 0.117 | 0.213 | 0.0 | **0.0*** | 2.143 | 0.003 |
| Iran (IROV) | Ovis vignei | 0.356 | 0.273 | 0.083 | 0.0 | **0.0*** | 0.848 | 0.002 |
| Various (ISGC) | Ovis aries | 0.228 | 0.105 | 0.122 | 0.0 | **0.0*** | 1.214 | 0.003 |
| Morocco (MOOA) | Ovis aries | 0.225 | 0.090 | 0.135 | 0.0 | **0.0*** | 1.331 | 0.002 |
| Barbados | Chlorocebus sabaeus | 0.368 | 0.109 | 0.260 | 0.0 | **0.0*** | 2.596 | 0.001 |
| Central African Republic (CAR) | Chlorocebus sabaeus | 0.256 | 0.073 | 0.183 | 0.0 | **0.0*** | 1.827 | 0.002 |
| Ethiopia | Chlorocebus sabaeus | 0.135 | 0.047 | 0.088 | 0.002 | **0.020*** | 0.885 | 0.002 |
| Gambia | Chlorocebus sabaeus | 0.198 | 0.043 | 0.154 | 0.0 | **0.0*** | 1.533 | 0.002 |
| Kenya | Chlorocebus sabaeus | 0.378 | 0.253 | 0.125 | 0.0 | **0.0*** | 1.278 | 0.001 |
| Nevis | Chlorocebus sabaeus | 0.055 | 0.035 | 0.020 | 0.136 | 0.618 | 0.206 | 0.001 |
| South Africa (SA) | Chlorocebus sabaeus | 0.227 | 0.056 | 0.172 | 0.0 | **0.0*** | 1.704 | 0.002 |
| Saint Kitts (SK) | Chlorocebus sabaeus | 0.061 | 0.035 | 0.026 | 0.108 | 0.618 | 0.265 | 0.001 |
| Zambia | Chlorocebus sabaeus | 0.175 | 0.073 | 0.102 | 0.007 | 0.056 | 1.016 | 0.002 |
| African (AFR) | Homo sapiens | 0.080 | 0.003 | 0.077 | 0.035 | 0.245 | 0.747 | 0.00071 |
| Ad Mixed American (AMR) | Homo sapiens | 0.041 | −0.003 | 0.045 | 0.119 | 0.618 | 0.435 | 0.00056 |
| East Asian (EAS) | Homo sapiens | 0.058 | 0.007 | 0.051 | 0.142 | 0.618 | 0.496 | 0.00051 |
| European (EUR) | Homo sapiens | 0.026 | 0.014 | 0.011 | 0.289 | 0.618 | 0.109 | 0.00054 |
| South Asian (SAS) | Homo sapiens | 0.085 | 0.008 | 0.076 | 0.103 | 0.618 | 0.744 | 0.00056 |

##### Mutation-selection codon model at site level ($\alpha$=0.025)

Figure S17:

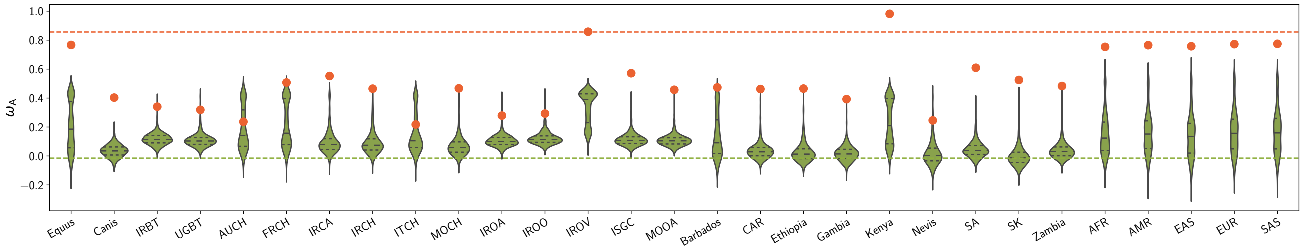


Table S18:

| Population | Species | $\omega_A$ Adaptive | $\langle\omega_A\rangle$ Nearly-neutral | $\Delta\omega_A$ | $p_v$ | $p_v^{adj}$ | $\frac{\Delta\omega_A}{\omega_A^{phy}}$ | $\pi_S$ |
|---|---|---|---|---|---|---|---|---|
| Diverse (Equus) | Equus caballus | 0.767 | 0.202 | 0.565 | 0.0 | **0.0*** | 0.650 | 0.00093 |
| Diverse (Canis) | Canis familiaris | 0.403 | 0.035 | 0.368 | 0.0 | **0.0*** | 0.420 | 0.001 |
| Iran (IRBT) | Bos taurus | 0.340 | 0.117 | 0.223 | 0.002 | **0.014*** | 0.257 | 0.003 |
| Uganda (UGBT) | Bos taurus | 0.319 | 0.110 | 0.208 | 0.011 | 0.050 | 0.240 | 0.003 |
| Australia (AUCH) | Capra hircus | 0.236 | 0.188 | 0.049 | 0.352 | 0.498 | 0.056 | 0.00099 |
| France (FRCH) | Capra hircus | 0.507 | 0.213 | 0.295 | 0.0 | **0.0*** | 0.337 | 0.00097 |
| Iran (IRCA) | Capra aegagrus | 0.552 | 0.096 | 0.456 | 0.0 | **0.0*** | 0.523 | 0.001 |
| Iran (IRCH) | Capra hircus | 0.465 | 0.092 | 0.373 | 0.0 | **0.0*** | 0.427 | 0.001 |
| Italy (ITCH) | Capra hircus | 0.217 | 0.149 | 0.068 | 0.249 | 0.498 | 0.078 | 0.001 |
| Morocco (MOCH) | Capra hircus | 0.467 | 0.074 | 0.393 | 0.0 | **0.0*** | 0.450 | 0.001 |
| Iran (IROA) | Ovis aries | 0.279 | 0.107 | 0.172 | 0.010 | 0.050 | 0.197 | 0.002 |
| Iran (IROO) | Ovis orientalis | 0.293 | 0.123 | 0.170 | 0.008 | **0.048*** | 0.195 | 0.003 |
| Iran (IROV) | Ovis vignei | 0.858 | 0.334 | 0.524 | 0.0 | **0.0*** | 0.603 | 0.002 |
| Various (ISGC) | Ovis aries | 0.572 | 0.114 | 0.457 | 0.0 | **0.0*** | 0.523 | 0.003 |
| Morocco (MOOA) | Ovis aries | 0.457 | 0.108 | 0.349 | 0.0 | **0.0*** | 0.399 | 0.002 |
| Barbados | Chlorocebus sabaeus | 0.474 | 0.137 | 0.337 | 0.0 | **0.0*** | 0.388 | 0.001 |
| Central African Republic (CAR) | Chlorocebus sabaeus | 0.462 | 0.037 | 0.426 | 0.0 | **0.0*** | 0.490 | 0.002 |
| Ethiopia | Chlorocebus sabaeus | 0.465 | 0.025 | 0.440 | 0.0 | **0.0*** | 0.507 | 0.002 |
| Gambia | Chlorocebus sabaeus | 0.392 | 0.019 | 0.373 | 0.0 | **0.0*** | 0.430 | 0.002 |
| Kenya | Chlorocebus sabaeus | 0.982 | 0.229 | 0.753 | 0.0 | **0.0*** | 0.871 | 0.001 |
| Nevis | Chlorocebus sabaeus | 0.246 | 0.021 | 0.225 | 0.034 | 0.102 | 0.260 | 0.001 |
| South Africa (SA) | Chlorocebus sabaeus | 0.609 | 0.048 | 0.562 | 0.0 | **0.0*** | 0.645 | 0.002 |
| Saint Kitts (SK) | Chlorocebus sabaeus | 0.525 | 0.001 | 0.524 | 0.0 | **0.0*** | 0.603 | 0.001 |
| Zambia | Chlorocebus sabaeus | 0.483 | 0.043 | 0.440 | 0.0 | **0.0*** | 0.507 | 0.002 |
| African (AFR) | Homo sapiens | 0.753 | 0.151 | 0.602 | 0.0 | **0.0*** | 0.690 | 0.00071 |
| Ad Mixed American (AMR) | Homo sapiens | 0.765 | 0.162 | 0.603 | 0.0 | **0.0*** | 0.691 | 0.00056 |
| East Asian (EAS) | Homo sapiens | 0.758 | 0.140 | 0.618 | 0.0 | **0.0*** | 0.709 | 0.00051 |
| European (EUR) | Homo sapiens | 0.772 | 0.165 | 0.607 | 0.0 | **0.0*** | 0.696 | 0.00054 |
| South Asian (SAS) | Homo sapiens | 0.775 | 0.168 | 0.607 | 0.0 | **0.0*** | 0.697 | 0.00056 |

- At the gene level, $\omega_A^{polyDFE}$ for nearly-neutral genes is the range $[-0.01, 0.27]$, while $\omega_A^{polyDFE}$ for genes under adaptation (at the phylogenetic scale) is in the range $[0.04, 42]$.
- At the site level, $\omega_A^{polyDFE}$ for nearly-neutral sites is the range $[0.0, 0.34]$, while $\omega_A^{polyDFE}$ for sites under adaptation (at the phylogenetic scale) is in the range $[0.40, 0.98]$.

The estimation of $\omega_A^{polyDFE}$ computed with polyDFE using polymorphism and divergence data with either a the model C (continuous DFE) or D (discrete DFE) are quite different in absolute value. The underlying assumption for the mathematical constrains on the DFE thus have a large impact on the estimation of $\omega_A^{polyDFE}$ while the true underlying DFE is unknown.
